# A genome-wide mutational constraint map quantified from variation in 76,156 human genomes

**DOI:** 10.1101/2022.03.20.485034

**Authors:** Siwei Chen, Laurent C. Francioli, Julia K. Goodrich, Ryan L. Collins, Masahiro Kanai, Qingbo Wang, Jessica Alföldi, Nicholas A. Watts, Christopher Vittal, Laura D. Gauthier, Timothy Poterba, Michael W. Wilson, Yekaterina Tarasova, William Phu, Mary T. Yohannes, Zan Koenig, Yossi Farjoun, Eric Banks, Stacey Donnelly, Stacey Gabriel, Namrata Gupta, Steven Ferriera, Charlotte Tolonen, Sam Novod, Louis Bergelson, David Roazen, Valentin Ruano-Rubio, Miguel Covarrubias, Christopher Llanwarne, Nikelle Petrillo, Gordon Wade, Thibault Jeandet, Ruchi Munshi, Kathleen Tibbetts, gnomAD Project Consortium, Anne O’Donnell-Luria, Matthew Solomonson, Cotton Seed, Alicia R. Martin, Michael E. Talkowski, Heidi L. Rehm, Mark J. Daly, Grace Tiao, Benjamin M. Neale, Daniel G. MacArthur, Konrad J. Karczewski

## Abstract

The depletion of disruptive variation caused by purifying natural selection (constraint) has been widely used to investigate protein-coding genes underlying human disorders, but attempts to assess constraint for non-protein-coding regions have proven more difficult. Here we aggregate, process, and release a dataset of 76,156 human genomes from the Genome Aggregation Database (gnomAD), the largest public open-access human genome reference dataset, and use this dataset to build a mutational constraint map for the whole genome. We present a refined mutational model that incorporates local sequence context and regional genomic features to detect depletions of variation across the genome. As expected, proteincoding sequences overall are under stronger constraint than non-coding regions. Within the non-coding genome, constrained regions are enriched for known regulatory elements and variants implicated in complex human diseases and traits, facilitating the triangulation of biological annotation, disease association, and natural selection to non-coding DNA analysis. More constrained regulatory elements tend to regulate more constrained protein-coding genes, while non-coding constraint captures additional functional information underrecognized by gene constraint metrics. We demonstrate that this genome-wide constraint map provides an effective approach for characterizing the non-coding genome and improving the identification and interpretation of functional human genetic variation.

## Introduction

The expansion in the scale of human whole-genome or exome reference data has allowed characterization of the patterns of variation in human genes. With these data it is possible to directly assess the strength of negative selection on loss-of-function (LoF) and missense variation by modeling “constraint,” the depletion of variation in a gene compared to an expectation conditioned on that gene’s mutability. Using coding variant data from sequencing of more than 125K humans^1^, we previously developed a constraint metric that classifies each protein-coding gene along a spectrum of LoF intolerance^1^, providing a valuable resource for studying the functional significance of human genes^2–5^. Although of outsized biological importance, protein-coding regions comprise less than 2% of the human genome, and the vast non-coding genome has been much less characterized, even though the importance of non-coding variation in human complex diseases has been long recognized^6–10^.

Several challenges arise when extending the gene constraint model to the non-coding space. First, the sample size of human whole-genome reference data has been relatively small compared to the exome, limiting the power of detecting depletions of variation at a fine scale. Second, our detailed understanding of coding region exon structure and effect of specific variants on amino acid translation enables a precision not available in non-coding analysis. Third, there is a strong expectation from Mendelian genetics and existing constraint analyses that the coding regions, while a small fraction of the genome, harbor a massively disproportionate amount of rare and common disease mutations under selection. Fourth, the mutation rate in non-coding regions is highly heterogeneous, and can be affected not only by local sequence context as commonly modeled in gene constraint metrics but also by a variety of genomic features at larger scales^11,12^.

Current methods attempting to evaluate non-coding constraint can be broadly divided into three categories: 1) context-dependent mutational models that assess the deviation of observed variation from an expectation based on the sequence composition of *k*-mers (e.g., Orion^13^, CDTS^14^, DR^15^); 2) machinelearning classifiers that are trained to differentiate between disease-associated variants and benign variants (e.g., CADD^16^, GWAVA^17^, JARVIS^18^); and 3) phylogenetic conservation scores that use comparative genomics data to infer evolutionary constraint (e.g., phastCons^19^, phyloP^20^). While all these methods aid in our understanding of the non-coding genome, each suffer from limitations/biases, respectively as 1) overlooking the influence of regional genomic features beyond the scale of flanking nucleotides on mutation rate; 2) a strong dependence on the availability of well-characterized functional mutations as training data; and 3) compromised power to detect regions that have only recently been under selection in the human lineage and may have a functional impact on human-specific traits or diseases.

Here we present a genome-wide map of human constraint, generated from a high-quality set of variant calls from 76,156 whole-genome sequences (gnomAD v3.1.2 https://gnomad.broadinstitute.org). We describe an improved model of human mutation rates that jointly analyzes local sequence context and regional genomic features and quantifies the depletion of variation in tiled windows across the entire genome. By building a more comprehensive picture of genic constraint rather than solely focusing on coding variation, we facilitate the functional interpretation of non-coding regions and improve the characterization of gene function in the context of the regulatory network. Our study aims to depict a genome-wide view of how natural selection shapes patterns of human genetic variation and generate a more comprehensive catalog of functional genomic elements with potential clinical significance.

## Results

### Aggregation and quality control of genome sequence data

We aggregated, reprocessed, and performed joint variant-calling on 153,030 whole genomes mapped to human genome reference build GRCh38, of which 76,156 samples were retained as high-quality sequences from unrelated individuals, without known severe pediatric disease, and with appropriate consent and data use permissions for the sharing of aggregate variant data. Among these samples, 36,811 (48.3%) are of non-European ancestry, including 20,744 individuals with African ancestries and 7,647 individuals with admixed Amerindigineous ancestries. After stringent quality control (see Supplementary Information), we discovered a set of 644,267,978 high-confidence short nuclear variants (single nucleotide/indel variants; gnomAD v3.1.2), of which 390,393,900 low-frequency (allele frequency [AF]≤1%), high-quality single nucleotide variants were used for building the genome-wide constraint map. These correspond to approximately one variant every 4.9 bp (one low-frequency variant every 8 bp) of the genome, providing a high density of variation.

### Quantifying mutational constraint across the genome

To construct a genome-wide mutational constraint map, we divided the genome into continuous non-overlapping 1kb windows, and quantified constraint for each window by comparing the expected and the observed variation in our gnomAD dataset. Here, we implemented a refined mutational model, which incorporates trinucleotide sequence context, base-level methylation, and regional genomic features to predict expected levels of variation under neutrality. In brief, we estimated the relative mutability for each single nucleotide substitution with one base of adjacent nucleotide context (e.g., ACG -> ATG), with adjustment for the effect of methylation on mutation rate at CpG sites, which become saturated for mutation at sample sizes above ~10K genomes^21^ (Extended Fig. 1a,b; Methods). Meanwhile, we adjusted the effects of regional genomic features for each trinucleotide mutation rate based on the occurrence of *de novo* mutations (*N*=413,304 previously detected in family-based whole-genome sequencing studies^22,23^; Extended Fig. 1c), and then applied it to establish the expected number of variants per 1kb across the entire genome (Methods).

We quantified the deviation from expectation for each 1kb window using a Z score^24^ (Methods; Extended Fig. 1d,e), which was centered around zero for non-coding regions (median=0.08), and was significantly higher (more constrained) for windows containing any protein-coding sequences (median=1.47, Wilcoxon *P*<10^−200^; **Fig. 1a**). The constraint Z score is positively correlated with the percentage of coding bases in a window and presented a substantial shift towards higher constraint for exonic sequences from directly concatenating coding exons into 1kb windows (median=3.17; Extended Fig. 2a-c). About 3.12% and 0.05% of the non-coding windows exhibited constraint as strong as the 50^th^ and 90^th^ percentile of exonic regions (Extended Fig. 2d). Comparing our Z score against the adjusted proportion of singletons (APS) score, a measure of constraint developed for structural variation (SV)^25^, we found a significant correlation (linear regression beta=0.01, *P*=4.3×10^−65^, **Fig. 1b**; Methods), providing an internal validation of our approach.

**Fig. 1:**
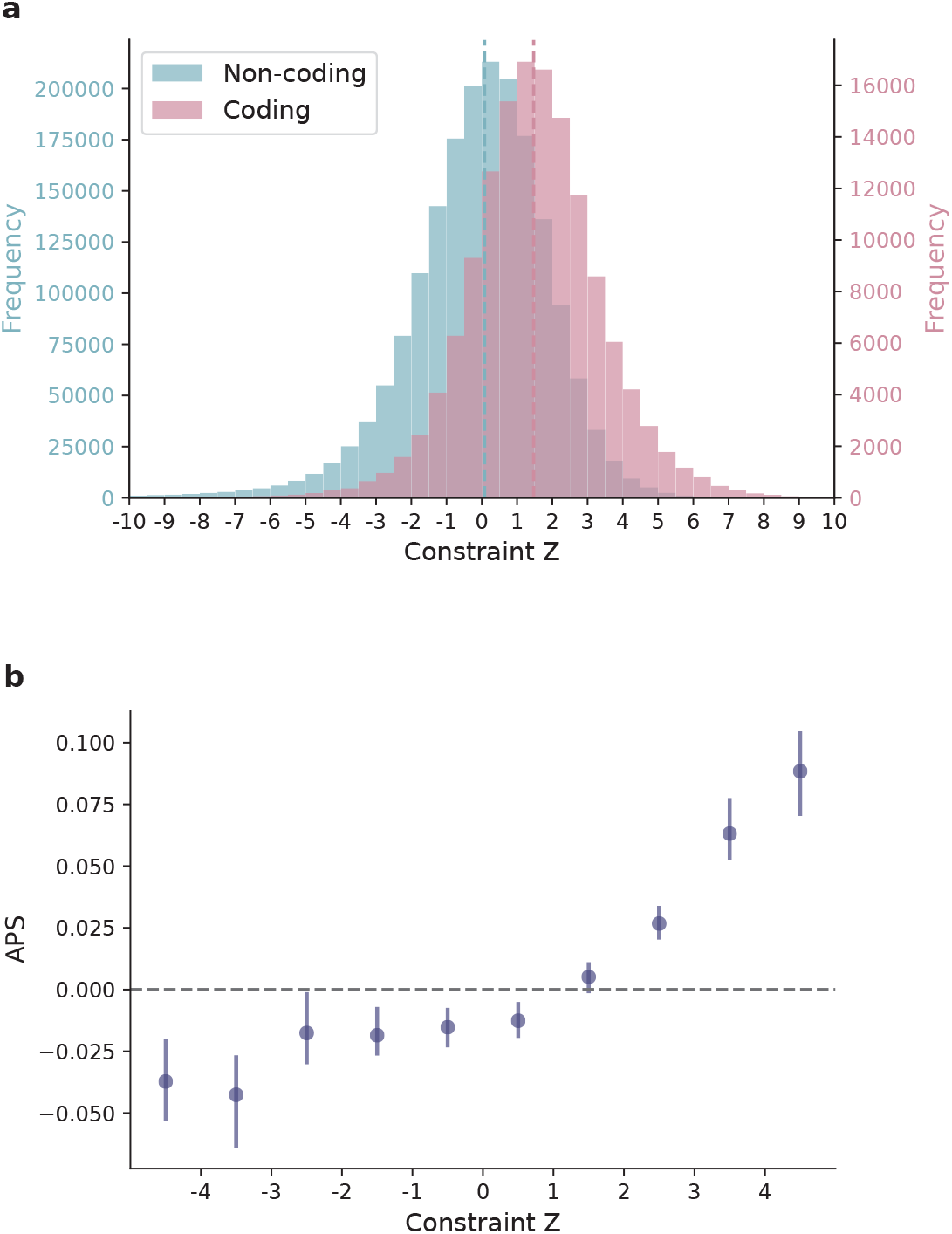
Distribution of constraint Z scores across the genome. **a**, Histograms of constraint Z scores for 1,984,900 1kb windows across the human autosomes. Windows overlapping coding regions (N=141,341 with ≥ 1bp coding sequence; red) overall exhibit a higher constraint Z (stronger negative selection) than windows that are exclusively non-coding (N=1,843,559; blue); dashed lines indicate the medians. **b**, The correlation between constraint Z score and the adjusted proportion of singletons (APS) score developed for structural variation (SV) constraint. A collection of 116,184 autosomal SVs were assessed using constraint Z score by assigning each SV the highest Z among all overlapping 1kb windows, which shows a signficant positive correlation with the SV constraint metric APS. Error bars indicate 100-fold bootstrapped 95% confidence intervals of the mean values.

### Investigating genomic properties of non-coding regions under constraint

To further validate our constraint metric and investigate the functional relevance of non-coding regions under selection, we examined the correlation between our constraint Z score and several annotations of functional non-coding sequences (**Fig. 2a**). First, we found that candidate cis-regulatory elements (cCREs, derived from ENCODE^26^ integrated DNase- and ChIP-seq data) are significantly enriched in the most constrained percentile of the genome (Z≥4, OR=2.77 compared to the genome-wide average, Fisher’s exact *P*<10^−200^); cCREs with a promoter-like signature (cCRE-PLS) presented the strongest enrichment (OR=7.28), followed by elements with a proximal/distal enhancer-like signature (pELS OR=4.35, dELS OR=2.14), and as a negative control, elements bound by CTCF but not associated with a regulatory signature showed no enrichment (CTCF-only OR=0.82). These patterns indicate that a large fraction of the constrained non-coding regions may serve a regulatory role, in line with previous findings^13,14,18^. Similarly, significant enrichment was found for an independent set of active, *in vivo*-transcribed enhancers (identified by FANTOM CAGE analyses^27^; OR=3.58) and super enhancers^28^ (OR=3.41), which are groups of enhancers in close genomic proximity regulating genes important for cell type specification^29^. By aggregating the regulatory annotations, we estimated that ~10.4% and ~6.3% of promoters and enhancers, respectively, are under selection as strong as coding exons on average (Extended Fig. 3a; Methods). A much higher proportion, 22.2%, was found for sequences encoding microRNAs (miRNAs), which are increasingly recognized as key mediators in various developmental and physiological processes^30^. In contrast, only 3.7% of long non-coding RNAs (lncRNAs) exhibited such strong constraint, similar to that of non-coding regions overall (3.1%; Extended Fig. 2d and 3b).

**Fig. 2:**
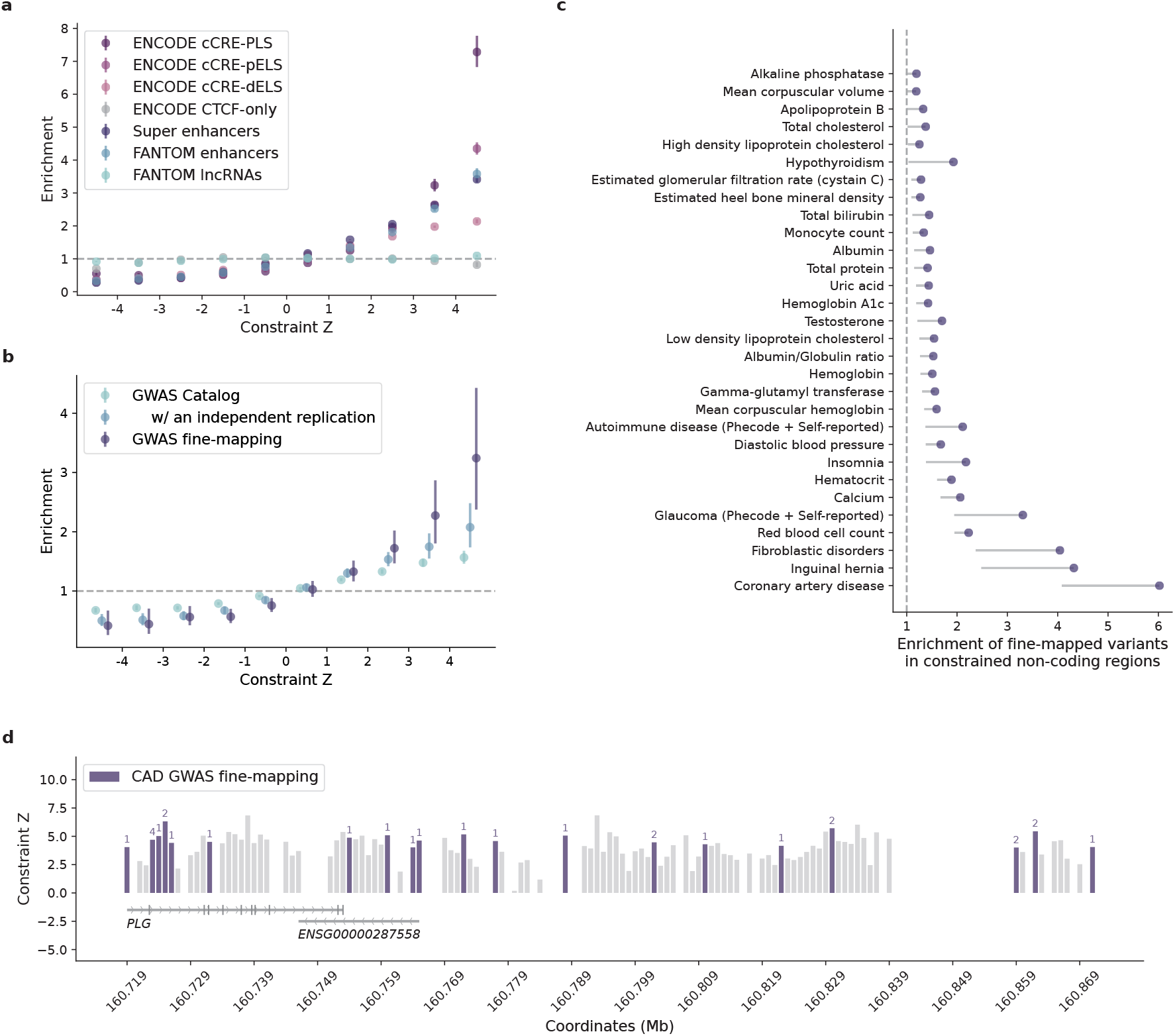
Correlation between constraint Z score and functional non-coding annotations. **a**,**b,** Distributions of candidate regulatory elements (**a**) and GWAS variants (**b**) along the spectrum of non-coding constraint. Enrichment was evaluated by comparing the proportion of non-coding 1kb windows, binned by constraint Z, that overlap with a given functional annotation to the genome-wide average. Error bars indicate 95% confidence intervals of the odds ratios. **c**, Enrichment of fine-mapped variants in constrained non-coding regions (Z≥4). Credible set (CS)-trat pairs with a significant enrichment are shown, ordered by the lower bound of 95% confidence interval; only lower bounds are shown for presentation purposes. **d**, The distribution of variants fine-mapped for coronary artery disease (CAD) in constrained regions (Z≥4) of *PLG.* Each bar shows the constraint Z score of a 1kb window (gaps indicate windows removed by quality filters); windows containing fine-mapped variants are colored by purple, and the number of variants in each window is annotated on top of the bar correspondingly. Ten variants are located within *PLG* introns, four are mapped to the antisense gene of *PLG* (ENSG00000287558), and 14 reside in the downstream intergenic regions.

We next examined the distribution of potentially functional non-coding variants on the constraint spectrum. There was significant enrichment for non-coding variants implicated by genome-wide association studies (GWAS) in the constrained end of the genome: 837/19,471 constrained windows [Z≥4] overlapped with GWAS Catalog^31^ annotations (OR=1.57 compared to the genome-wide average of 51,430/1,843,559, Fisher’s exact *P*=2.5×10^−32^, **Fig. 2b**; Methods). The enrichment became stronger when restricted to the subset of variants that had been replicated by an independent study (OR=2.08, *P*=4.1×10^−13^). Moreover, further strong signals were found for likely causal GWAS variants fine-mapped for 148 complex diseases and traits in large-scale biobanks^32^ (OR=3.24, *P*=3.0×10^−10^; Methods). Across the 95% credible set (CS)-trait pairs, strong enrichment was predominantly seen in disease phenotypes, including coronary artery disease (CAD), inguinal hernia, fibroblastic disorders, and glaucoma (ORs 3.31-6.02, **Fig. 2c**; Methods). In the 95% CS of CAD, for instance, the highest constraint score was found for rs1897107 and rs1897109 (both within the same genomic window chr6:160725000-160726000, Z=6.32); high constraint (Z≥4) was also found for 26 variants from the same CS (totaling 28/52), which together spanned a ~153 kb sequence downstream of the gene *PLG* (**Fig. 2d**). *PLG* encodes the plasminogen protein that circulates in blood plasma and is converted to plasmin to dissolve the fibrin of blood clots. While dysregulation of the PLG-plasmin system has been frequently associated with CAD^33–38^, no specific variants in *PLG* have been implicated. Our results prioritized a set of non-coding variants in highly constrained regions of *PLG,* which adds quantitative evidence to the implication of *PLG* in CAD and may help direct or prioritize follow-up functional experiments.

Collectively, these results demonstrated a significant positive correlation between constraint and existing functional non-coding annotations, validating our non-coding constraint metric. Yet, we suggest that our metric provides additional information for the characterization of non-coding regions. For instance, prioritizing ENCODE cCREs by constraint Z score revealed increasingly stronger GWAS enrichment in the more constrained cCREs (Extended Fig. 4a), and constrained regions outside cCREs also captured significant signals, reflecting the value of non-coding constraint independent of regulatory annotations. Moreover, besides prioritizing existing GWAS results, constraint can be used as a prior for statistical fine-mapping. Using UK Biobank (UKBB) traits as examples, incorporating constraint Z score into the functionally informed fine-mapping model^39^ predicted ~13K variant-trait pairs to have an increased posterior inclusion probability of causality (△PIP≥0.01), in which 164 likely causal associations were newly identified at PIP≥0.8 (Extended Fig. 4b; Methods). While only functional tests can ultimately validate the underlying causality, our constraint map presents a valuable resource for expanding or refining the catalog of functional non-coding variants in the human genome.

### Comparing constraint Z score to other genome-wide predictive scores

To benchmark the performance of our constraint map in prioritizing non-coding variants, we extended the analyses of GWAS variants to compare constraint Z score to other population genetics-based constraint metrics (Orion^13^, CDTS^14^, gwRVIS^18^, and DR^15^). Specifically, we assessed the performance of different metrics in identifying putative functional non-coding variants – as aforementioned, a) GWAS Catalog^31^ variants (N=9,229 with an independent replication); b) GWAS fine-mapping^32^ variants (N=2,191), and additionally, c) a subset of high-confidence causal variants from b (N=140); and d) pathogenic Mendelian variants (N=288 from ClinVar^40^) – against background variants in the population with a similar allele frequency (hereafter referred to as “positive” and “negative” variant set, respectively; Methods). Overall, constraint Z score achieved the highest performance across all comparisons, as measured by the area under curve (AUC) statistic (**Fig. 3a,b** and Extended Fig. 5). The performance was also more stable than others when varying the allele frequency threshold for the negative variant set (Extended Fig. 5). This may be due to other metrics being informed by the site frequency spectrum, which made the classification performance sensitive to differences in allele frequency between the positive and negative variants. We also showed that our performance was robust to the artificial break of genomic windows (non-overlapping 1kb) by reconstructing constraint Z scores in a sliding-window (1kb stepped by 100bp) approach as adopted by other metrics (Extended Fig. 6).

**Fig. 3:**
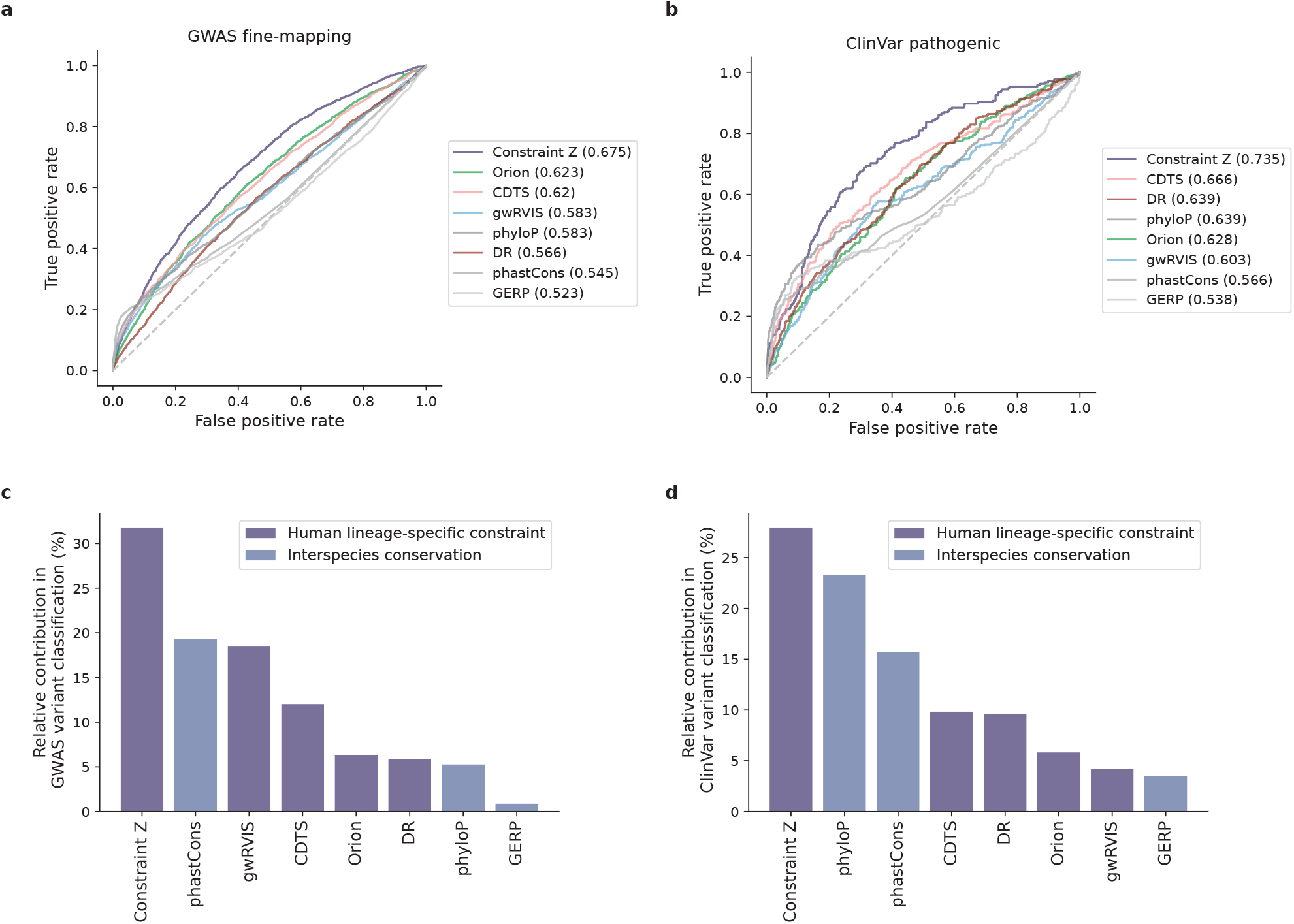
Performance of constraint Z score and other predictive scores in prioritizing non-coding variants. **a**,**b**, Receiver operating characteristic (ROC) curves of constraint Z score and other seven metrics in classifying putative functional non-coding variants – 2,191 GWAS fine-mapping variants (**a**) and 288 ClinVar pathogenic variants (**b**) – against background variants in the population. The performance of each metric was measured and ranked by the area under curve (AUC) statistic. **c**,**d**, The relative contribution of different metrics in classifying GWAS variants (**b**) and ClinVar variants (**c**). The eight metrics were modeled as eight independent predictors for the classification, and the relative contribution of one predictor over another was evaluated by estimating their additional *R*^2^contributions across all subset models.

Extending the comparison to include phylogeny-based conservation scores (phyloP^20^, phastCons^19^, and GERP^41^) revealed relatively low performance compared to the population genetics-based constraint metrics (**Fig. 3a,b**). The conservation scores were weakly correlated with constraint (Spearman’s rank correlation coefficient 0.017-0.19, Extended Fig. 7), suggesting that intraspecies (human lineage-specific) constrained regions complement, rather than being a subset of, regions that are conserved across species. Each individual metric also contributed to the classification when modeled as independent predictive variables (**Fig. 3c,d**; Methods), reinforcing the complementary nature of different approaches. Variants that were uniquely captured by constraint Z score, for instance, tended to be in regions with high recombination rates (3.45-fold the rest of the positive variant set) and high DNA methylation (2.74-fold; Methods), both associated with an increased mutation rate that had been adjusted in our refined mutational model. To further illustrate this improvement, we rebuilt our constraint model from solely the local sequence context, i.e., without adjustment on mutation rate by regional genomic features, and confirmed that constraint Z score outperformed such metrics (Extended Fig. 6). Altogether, we demonstrate that constraint Z score is an effective metric for identifying functional variants in the noncoding genome; at the same time, we suggest that a combination of different metrics is likely to provide the most informative results.

### Exploring non-coding dosage sensitivity in the constrained genome

Besides single nucleotide variants (SNVs) that have been extensively studied in GWAS, copy number variants (CNVs) causing dosage alterations (deletions/loss or duplications/gain) of DNA represent another significant risk factor for human disease^42–47^. Yet, unlike SNVs, CNVs can be large and determining the “minimal critical region”^48^ with a pathogenic effect has been a major challenge. Although CNVs primarily affect non-coding sequences, the most commonly studied mechanism is still the dosage alteration of overlapping protein-coding genes^49^. Using our genome-wide constraint map, we explored the possibility that constrained non-coding regions are also sensitive to a dosage effect, which may underlie the pathogenicity of corresponding CNVs.

We surveyed a collection of ~100K CNVs from a genome-wide CNV morbidity map of developmental delay and congenital birth defects^50,51^. There was a substantial excess of CNVs that affected constrained noncoding regions (Z≥4) among individuals with developmental disorders (DD cases) in comparison to healthy controls (42.6% versus 12.5%, OR=5.21, Fisher’s exact *P*<10^−200^, **Fig. 4a**; Methods). Moreover, of the 19 loci that had been previously identified as pathogenic^50^, all but one (94.7%) affected constrained non-coding regions; the high incidence was recapitulated in a curated set of ~4K putative pathogenic CNVs (85.5% in ClinVar^40^, **Fig. 4a**). Importantly, the case-control enrichment remained significant, albeit attenuated, after adjusting for the size and gene content of each CNV and when being tested in the subset of CNVs that are exclusively non-coding (**Fig. 4b**; Methods). Non-coding constraint presented high association with DD CNVs conditioning on gene constraint (log[OR]=1.06, logistic regression *P*<10^−100^), lending support to the possibility that dosage alteration of constrained non-coding regions may be an alternative explanation for the mechanism of CNVs underlying DDs.

**Fig. 4:**
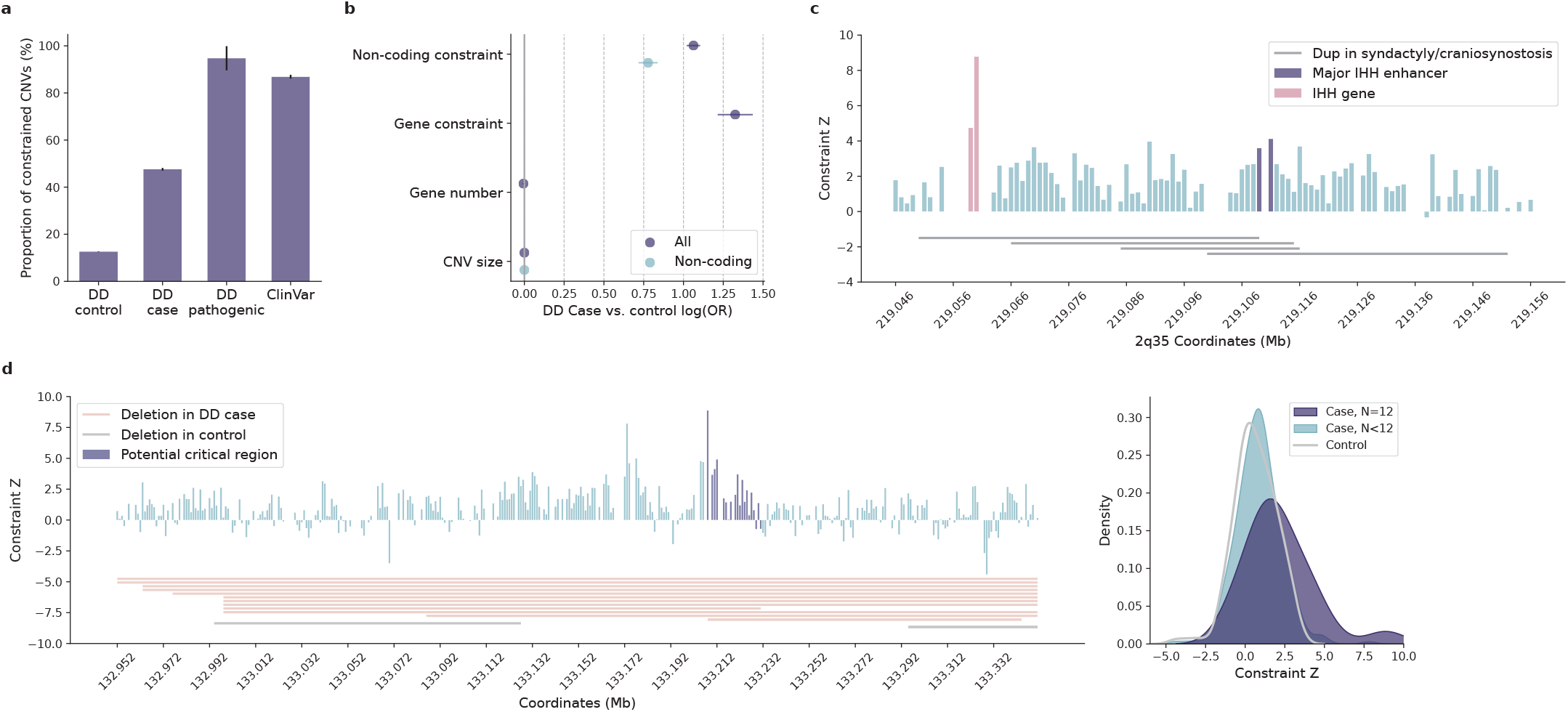
Contribution of non-coding constraint in evaluating copy number variants (CNVs). **a**, Proportions of constrained CNVs (Z≥4) identified in individuals with developmental delay (DD cases) versus healthy controls. Constrained CNVs are more common in DD cases than controls (7,239/17,004=42.6% versus 10,403/83,526=12.5%) and are most frequent for CNVs previously implicated as pathogenic (18/19=94.7% by DD and 3,433/4,014=85.5% by ClinVar). **b**, Contribution of non-coding constraint to predicting CNVs in DD cases versus controls. Non-coding constraint remains a significant predictor for the case/control status of CNVs after adjusting for gene constraint (LOEUF score), gene number, and size of CNVs (purple), as well as being tested in the subset of non-coding CNVs (blue). Error bars indicate 95% confidence intervals of the log odds ratios. **c**, CNVs at the *IHH* locus associated with synpolydactyly and craniosynostosis. The four implicated duplications (grey horizontal bars) span a ~102kb sequence upstream of *IHH.* Each vertical bar shows the constraint Z score of a 1kb window within the locus, with the highest score overlapping the *IHH* gene (red) and the highest non-coding score overlapping the major *IHH* enhancers (purple); gaps indicate windows removed by quality filters. **d**, Non-coding CNVs with the highest constraint Z score identified in DD cases. The highest-scored window is located within the potential “critical region”^48^ (purple vertical bars) shared by 12 DD deletions (red horizontal bars; grey indicates two deletions observed in controls). The critical region overall, has a significantly higher constraint Z score than the other regions affected by DD or control deletions, as shown in the kernel density estimate (KDE) plot on the right.

One known example of pathogenic non-coding dosage alteration is the duplication of *IHH* regulatory domain in synpolydactyly and craniosynostosis^52–54^. The four implicated duplications covered a ~102kb sequence upstream of *IHH,* with a ~10kb overlapping region (“critical region”^48^; **Fig. 4c**). The region contained no genes but exhibited high levels of constraint (median Z=2.52, Wilcoxon *P*=1.3×10^−3^ compared to the rest of the genome). The most constrained window (chr2:219111000-219112000, Z=4.12) overlapped with the major enhancer of *IHH,* the duplication of which has been shown to result in dosage-dependent *IHH* misexpression and consequently syndactyly and malformation of the skull^54^. This result highlights a potential use of the constraint metric to prioritize non-coding regions within large CNVs. As a further illustration, we examined a set of non-coding CNVs that had the highest constraint score among the DD cases. The most constrained genomic window (chr11:133208000-133209000, Z=8.87) was affected by 12 deletions spanning a ~400kb non-coding sequence (**Fig. 4d**). While of varying size, the deletions shared a common region of ~20kb (potential “critical region”), which encompassed the most constrained window and overall, showed a significantly higher constraint than the other affected regions (median Z = 1.63 vs 0.84, Wilcoxon *P*=1.6×10^−3^; **Fig. 4d**). In addition, the ~400kb sequence also harbored two deletions from healthy controls, which interestingly, overlapped with the two lowest constraint scores within the region and were significantly less constrained than those from DD cases (median Z = 1.07 vs 0.62, Wilcoxon *P*=4.74×10^−4^). These findings suggest that our constraint metric can be a useful indicator of critical regions affected by large CNVs, facilitating the identification of non-coding risk factors in CNV disease association studies.

### Leveraging non-coding constraint to improve gene function characterization

Given the significant role of constrained non-coding regions in gene regulation, it is natural to expect that more constrained regulatory elements would regulate more constrained genes. To test this, we analyzed the constraint for enhancers that had been linked to specific genes^55^ (Methods). More constrained noncoding regions were more frequently linked to regulating a gene (**Fig. 5a**), and as expected, enhancers linked to constrained genes (predicted by loss-of-function observed/expected upper bound fraction [LOEUF]^1^, or curated disease genes from ^56–58^; Methods) were significantly more constrained than those linked to presumably less constrained genes (median Z=2.71 versus 1.99, Wilcoxon *P*=1.3×10^−26^, **Fig. 5b**; Methods), thus supporting a correlated constraint between genes and their regulatory elements.

**Fig. 5:**
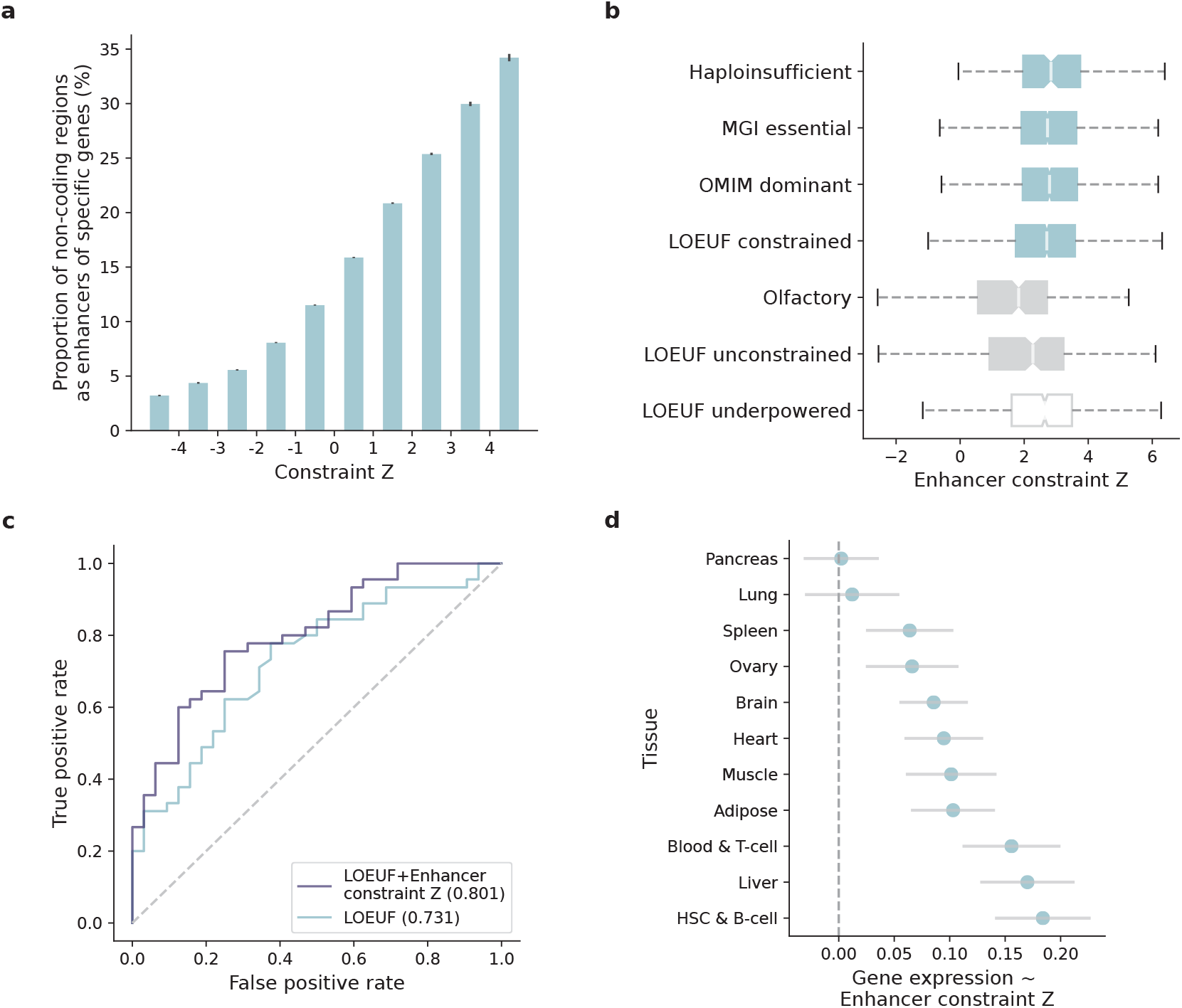
Correlation of constraint between non-coding regulatory elements and protein-coding genes. **a**, The proportion of non-coding 1kb windows overlapping with enhancers that were predicted to regulate specific genes, as a function of their constraint Z scores. More constrained non-coding regions are more frequently linked to a gene. Error bars indicate standard errors of the proportions. **b**, Comparison of the constraint Z scores of enhancers linked to constrained and unconstrained genes. Enhancers of established sets of constrained genes (four blue boxes) are more constrained than enhancers of presumably less constrained genes (two grey boxes). Enhancers of genes that are underpowered for gene constraint detection (“LOEUF underpowered”) present a higher constraint than those powered yet unconstrained genes (“LOUEF unconstrained”). **c**, Improvement of incorporating enhancer constraint into LOEUF in prioritizing underpowered genes. ROC curves and AUCs show the performance of two models using LOEUF (blue) and LOEUF+enhancer constraint Z score (purple) to classify constrained and unconstrained genes, tested on a set of 77 underpowered genes. **d**, Contribution of enhancer constraint to predicting gene expression in specific tissue types. The x-axis shows the linear regression coefficient of tissue-specific enhancer constraint predicting the expression level of target genes in matched tissue types, conditioning on gene constraint (LOEUF score). Error bars indicate 95% confidence intervals of the coefficient estimates.

On the other hand, a particularly interesting set of associations are the links between constrained enhancers and the “unconstrained” genes predicted by LOEUF, because these links may reflect functional significance of the “unconstrained” genes that had been previously unrecognized. The lack of predicted gene constraint can be explained by the design of LOEUF as a measure of intolerance to rare LoF variation, where small genes with few expected LoF variants are likely underpowered. Indeed, stratifying genes by the number of expected LoF variants showed a significantly higher enhancer constraint for genes that were underpowered (≤5 expected LoF variants)^1^ compared to genes that were sufficiently powered while scored as unconstrained (median Z=2.64 versus 2.27, Wilcoxon *P*=9.8×10^−4^, **Fig. 5a**). This suggests that certain underpowered genes may be functionally important but were not recognized in gene constraint evaluation. For instance, *ASCL2,* a basic helix-loop-helix (bHLH) transcription factor, had only 0.57 expected LoFs (versus 0 observed) across >125K exomes^1^; although being depleted for LoF variation, the absolute difference was too small to obtain a precise estimate of LoF intolerance. Yet, we found *ASCL2* had a highly constrained enhancer (Z=5.58), located ~16kb upstream of the gene, where >40% of the expected variants were depleted (188.6 expected versus 112 observed, chr11:2286000-2287000). The same genomic window also contained an eQTL chr11:2286192:G>T that was predicted to be significantly associated with *ASCL2* expression^59^; elevated *ASCL2* expression has been implicated in the development and progression of several human cancers^60–62^. This example highlights the value of non-coding constraint – as a complementary metric to gene constraint – for identifying functionally important genes.

A practical implementation of this finding is to integrate the constraint of regulatory elements into the modeling of gene constraint, which essentially borrows power from extending the functional unit of a gene to encompass its regulatory components. As a proof-of-principle, we tested whether adding the enhancer constraint Z score to LOEUF improves the prioritization of underpowered genes. The enhancer constraint was found a significant predictor of constrained genes (logistic regression *P*=7.4×10^−11^ conditioning on LOEUF) and significantly improved the performance of LOEUF in identifying constrained genes that were underpowered (AUC = 0.80 versus 0.73, bootstrap *P*=0.03, **Fig. 5b**; Methods). Moreover, such approaches would allow incorporation of tissue/cell-type specific information into gene constraint modeling given the diverse range of epigenomic data. We explored this by testing whether the constraint of tissue-specific enhancers is predictive of tissue-specific gene expression (as a proxy for tissue-specific gene function). The enhancer constraint Z score, again conditioning on LOEUF, was a significant predictor of the expression level of target genes in matched tissue types (**Fig. 5c**; Methods). These results further support the application of our constraint metric for improving the characterization of gene function. While we acknowledge that the biological consequences of mutations in enhancers are not clearly understood and thus natural selection may differ in strength depending on mechanistic consequence, an extended model to incorporate non-coding variation information in a biologically-informed way holds promise to facilitate our understanding of the molecular mechanisms underlying selection.

## Discussion

We have previously developed constraint metrics that leverage population-scale exome and genome sequencing data to evaluate genic intolerance to coding variation for each protein-coding gene^1,21^. Here, we adopted the same principle with an extended mutational model to assess constraint across the entire genome, using our latest release of gnomAD (v3.1.2), a dataset of harmonized high-quality whole-genome sequences from 76,156 individuals of diverse ancestries. Improvements to constraint modeling include unified fitting of the mutation rate for all substitution and trinucleotide contexts and inclusion of regional genomic features to refine the expected variation in non-coding regions (Methods). We validated our metric using a series of external functional annotations, with a focus on the non-coding genome, and demonstrated the value of our metric for prioritizing non-coding elements and identifying functionally important genes. We have made the constraint scores publicly accessible via the gnomAD browser (https://gnomad.broadinstitute.org).

One key challenge in quantifying non-coding constraint is the estimation of mutation rate under neutrality, which can be affected by various genomic features at different scales. To this end, we extended our previous mutational model, which computed the relative mutability of each substitution in a trinucleotide context, to include adjustment for the effects of regional genomic features. The adjustment was applied to each specific trinucleotide context and allowed a varying genomic scale for each specific feature (Methods). The added value of this adjustment was demonstrated by the improved performance of the constraint Z score in identifying functional variants (Extended Fig. 6). Our constraint metric also outperformed other genome-wide predictive scores, while each metric tended to provide complementary information. We note that all comparisons were restricted to non-coding regions for explicitly evaluating the metrics in prioritizing non-coding variants, and we further eliminated potential bias from nearby genes by recapitulating the results within regions >10kb away from any protein-coding exons (Supplementary Information). Overall, our constraint metric presented consistent, high performance in identifying functional non-coding variants in the human genome.

Despite the clear constraint signal identified for non-coding regions, many limitations exist. First, the lack of prior classification of the molecular consequences of non-coding variants, as analogous to “nonsynonymous” versus “synonymous” informed by the genetic code in coding regions, limits the resolution of non-coding constraint assessment (e.g., to measure constraint against “LoF” variation). While there are rich resources defining regulatory elements in the non-coding genome, no method is available for determining the impact of each possible variant on gene expression and the distribution of their effect sizes genome-wide. Further, the interpretation of non-coding constraint, especially in the context of gene regulation, can only be informative when considered in a particular context, such as a tissue/cell type, developmental stage, or environment. Such information is not inherently built into our constraint metric nor in the mutational dataset; thus *ad hoc* integration of external annotations (e.g., tissue-specific enhancers as analyzed in this study) is often necessary for justifying specific biological implications. Also, since the detection of depletion of variation is immune to negative selection after reproductive age, genomic regions involved in late-onset phenotypes are likely to go underrecognized.

Finally, while this is the largest dataset of human genomes examined to date for non-coding constraint, our method will substantially increase in power and resolution as sample sizes increase. Benchmarking on the depletion of variation seen in coding regions, we are currently well-powered to detect extreme noncoding constraint as strong as the 90^th^ percentile of coding exons of similar size, and we estimate a sample size of ~340K genomes to detect constraint as to the 50^th^ percentile (Extended Fig. 8a; Methods). Much larger sample sizes will be required for further increasing the resolution, for instance from 1kb to a 100bp scale, we would need ~5.3M samples (Extended Fig. 8b); under the current sample size, 1kb presented optimal performance when compared to a various window size tested from 100bp-3kb (Extended Fig. 8c). Meanwhile, we emphasize the importance of increasing ancestral diversity in population-scale datasets like gnomAD. A more diverse population would identify a larger number of rare variants, thereby increasing the power of detecting depletions of variation. We explicitly demonstrated this by reconstructing our constraint metric from the subset of European population and comparing it to that from an equal-sized subset containing all diverse populations – the latter was proven to achieve a higher predictive power (Extended Fig. 8d). Future efforts towards a larger, more diverse human reference dataset would empower finer studies of the influence of human demography on constraint metrics, facilitating a fuller understanding of the distribution and effect of human genetic variation.

Overall, our study demonstrates the value of the genome-wide constraint map in characterizing both noncoding regions and protein-coding genes, providing a significant step towards a comprehensive catalog of functional genomic elements for humans.

## Methods

### Aggregation, variant-calling, and quality control of gnomAD genome data

We aggregated whole genome sequence data from 153,030 individuals spanning projects from case-control consortia and population cohorts, in a similar fashion to previous efforts^1^. We harmonized these data using the GATK Best Practices pipeline and joint-called all samples using Hail^63^, and developed and utilized an updated pipeline of sample, variant, and genotype quality control to create a high-quality callset of 76,156 individuals, computing frequency information for several strata of this dataset based on attributes such as ancestry and sex for each of 644,267,978 short nuclear variants (see Supplementary Information).

### Estimation of trinucleotide context-specific mutation rates

We estimated the probability of a given nucleotide mutating to one of the three other possible bases in a trinucleotide context (XY1Z -> XY2Z), by computing the proportion of all possible variants observed per context in the human genome. Since CpG transitions begin to saturate (proportion observed approaching 1) at a sample size of ~10K genomes, we downsampled the gnomAD dataset to 1,000 genomes for this calculation. The computed proportion observed values, which represent the relative mutability of each trinucleotide context, were further scaled so that the weighted genome-wide average is the human perbase, per-generation mutation rate (1.2×10^−8^) to obtain the absolute mutation rates *μ.* To estimate the proportion of variants expected to be observed in the full gnomAD dataset of 76,156 genomes, we fitted the actual proportion observed in the dataset against *μ,* using an exponential regression that caps at 1 for refining the estimates of (near-)saturated variant types (R^2^=0.999, Extended Fig. 1a,b; Supplementary Data 1).

A total of 390,393,900 high-quality, rare (AF≤0.1%) variants observed in 76,156 gnomAD genomes, a dataset of 6,079,733,538 possible variants at 2,026,577,846 autosomal sites (30-32X coverage), were used in the calculation of trinucleotide context-specific mutation rates. The estimates are well-correlated with the mutation rates reported in previous independent studies and are highly stable across different AF thresholds in gnomAD (Supplementary Information).

### Adjustment of the effect of DNA methylation on CpG mutation rates

Given the strong effect of DNA methylation on increasing the mutation rate at CpG sites, we stratified all CpG sites by their methylation levels and computed the proportion observed within each context and methylation level. As an improvement to our previous methylation annotation (by averaging different tissues^1^), we analyzed methylation data from germ cells across 14 developmental stages, comprising eight from preimplantation embryos (sperm, oocyte, pronucleus, two-cell-, four-cell-, eight-cell-, morula-, and blastocyst-stage embryos)^64^ and six from primordial germ cells (7Wk, 10Wk, 11Wk, 13Wk, 17Wk, and 19Wk)^65^. For each stage, we computed methylation level at each CpG site as the proportion of wholegenome bisulfite sequencing reads corresponding to the methylated allele. To derive a composite score from the 14 stages, we regressed the observation of a CpG variant in gnomAD (0 or 1) on the methylation computed at the corresponding site (a vector of 14), and we used the coefficients from the regression model as weights to compute a composite methylation score for each CpG site. This metric was further discretized into 16 levels (by a minimum step of 0.05: [0,0.05], (0.05,0.1], (0.1,0.15], (0.15,0.2], (0.2,0.25], (0.25,0.3], (0.3,0.5], (0.5,0.55], (0.55,0.6], (0.6,0.65], (0.65,0.7], (0.7,0.75], (0.75,0.8], (0.8,0.85], (0.85,0.9], (0.9,1.0]) to stratify CpG variants in the mutation rate analysis.

### Adjustment of the effects of regional genomic features on mutation rates

To estimate the effects of regional genomic features on mutation rates under neutrality, we utilized *de novo* mutations (DNMs), as a proxy of spontaneous mutations, and fitted logistic regression models using the genomic features as predictive variables. A set of 413,304 unique DNMs were compiled from two large-scale family-based whole-genome sequencing studies^22,23^, and an exclusive set of 4,104,879 genomic sites (~10× the DNMs) randomly drew from the genome was used as the “nonmutated” background. For each DNM or background site, we computed 13 genomic features (see Collection of genomic features) at four scales by taking the mean value of 1kb, 10kb, 100kb, and 1Mb windows centering at the site. This generated a feature matrix of 13×4=52 columns and 413,304+4,104,879 =4,518,183 rows. The matrix was further divided based on the trinucleotide context of each DNM or background site (by row) to assess the effects of genomic features on context-specific mutation rates. In particular, for CpG contexts, features that were correlated with DNA methylation (GC content, CpG_island, short interspersed nuclear element, and nucleosome density), which had been used for adjusting CpG mutation rates, were excluded from the analysis.

For each trinucleotide context, we first performed univariable logistic regression to select features that are significantly associated with an increased/decreased probability of observing a DNM. Features with a significant association surpassing the Bonferroni correction for 13×4=52 tests were selected; if a feature was significant at multiple genomic scales, the smallest window size was selected for the highest resolution (Extended Fig. 1c). Next, we fitted multivariable logistic regression using the selected features to predict DNMs from the background. To control for multicollinearity, we transformed the input feature matrix using principal components analysis (PCA^66^) to generate decorrelated predictive variables (i.e., the principal components or PCs). The regression coefficients were the primary output of interest, which represent the effects of genomic features on increasing (a positive coefficient) or decreasing (a negative coefficient) the mutation rate, and were used for adjusting the expected number of variants in a given region. The selected features, the PCs, and the coefficients are summarized in Extended Fig. 1c and are available as pickle files for implementation (see Code availability in Supplementary Information).

### Prediction of expected number of variants per 1kb

Using the trinucleotide mutation rate estimates and the above adjustments, we computed the expected number of variants in a given 1kb genomic window as follow:

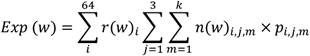

where *i* denotes one of the 64 trinucleotide contexts; *j* denotes one of the three bases substituting the central nucleotide; *m* denotes one of the *k* DNA methylation levels, where *k*=16 for CpG sites (see Adjustment of the effect of DNA methylation on CpG mutation rates) and *k*=1 for non-CpG sites (i.e., no stratification). Essentially, the expected value of variants in a genomic window *w* is calculated by multiplying the number of possible variants (*n*) in *w* by the probability of a variant (*p*) and summing across all trinucleotide contexts (*i*), substitutions (*j*), and methylation levels (*m*); *p_i,j,m_* is the trinucleotide mutation rate estimated in this study (as described in Estimation of trinucleotide context-specific mutation rates).

Additionally, *Exp* is adjusted by a factor *r,* which represents the effect of regional genomic features of *w* on mutation rate. For each *i*, specific features have been pre-selected and their effects on mutation rate have been estimated using logistic regression models (see Adjustment of the effects of regional genomic features on mutation rates). Denote the feature values, computed centering *w* and decorrelated by PCA, and the regression coefficients by ***x*** = {*x*_1_, *x*_2_,…, *x_t_*} and ***β*** = {*β*_1_, *β*_2_,.., *β_t_*}, respectively, where *t* is the number of selected features for i, the adjustment factor *r* is defined as the ratio of logit given *x*(*w*) to that of the genome-wide average 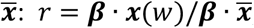; since the adjustment is specific to each trinucleotide context, *r* is further subscribed by *i*.

### Construction of constraint Z score

We created a signed Z score to quantify the depletion of variation (constraint) at a 1kb scale by comparing the observed variation to an expectation:

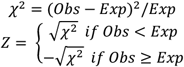

The observed variant count (*Obs*) is the number of unique rare (AF≤0.1%) variants in a 1kb window identified in the gnomAD dataset of 76,156 genomes, and the expected number of variants *(Exp)* is established as described above based on the sequence context and the regional genomic features of the 1kb window.

Constraint Z scores were created for 2,689,987 non-overlapping 1kb windows across the human genome, comprising 2,561,056 on autosomes and 128,931 on chromosome X. Due to the lack of DNM data on chromosome X, the genomic feature adjustment factor *r* was assessed using autosomal regions and extrapolated to chromosome X. We performed downstream analyses separately for autosomes and chromosome X and presented the former as primary, with the latter provided in Supplementary Information. For the analyses, we filtered the dataset to windows where 1) the sites contained at least 1,000 possible variants, 2) at least 80% of the observed variants passed all variant call filters (INFO/FILTER equals to “PASS”), and 3) the mean coverage in the gnomAD genomes was between 25-35X (or 20-25X for chromosome X). This resulted in 1,984,900 autosomal windows (77.5% of initial) for the primary analyses, of which 141,341 overlapped with coding regions and 1,843,559 were exclusively non-coding. The computed constraint Z scores are available in Supplementary Data 2. We also computed the sores in a sliding window approach (1kb stepped by 100bp) and provided them in Supplementary Data 3.

### Collection of genomic features

The 13 regional genomic features used for adjusting trinucleotide mutation rate are 1) GC content^67^, 2) low-complexity region^68^, 3) short and 4) long interspersed nuclear element^67^, distance from the 5) telomere and the 6) centromere^67^, 7) male and 8) female recombination rate^22^, 9) DNA methylation, 10) CpG island^67^, 11) nucleosome density^69^, 12) maternal and 13) paternal DNM cluster^70^. Data were downloaded from the referenced resources, lifted over to GRCh38 coordinates when needed using CrossMap^71^, and files in .bed or .BigWig format were processed using bedtools^72^ and bigWigAverageOverBed^69^ to obtain feature values within specific genomic windows.

### Correlation between constraint Z score and APS

As an internal validation, we compared our constraint Z score against the SV constraint score APS^25^. For each SV from the original study^25^, we assessed its constraint by assigning the highest Z score among all overlapping 1kb windows. The correlation between constraint Z and APS was evaluated across 116,184 high-quality autosomal SVs scored by both metrics, using a linear regression test. In Fig. 1b, the correlation was presented by the mean value of APS across ascending constraint Z score bins, with 95% confidence intervals computed from 100-fold bootstrapping.

### Correlation between constraint Z score and putative functional non-coding annotations

We validated the constraint metric using a number of external functional annotations, including 926,535 ENCODE cCREs^26^ (34,803 promoter-like [PLS], 141,830 proximal enhancer-like [pELS], 667,599 distal enhancer-like [dELS), and 56,766 CTCF-only elements), 63,285 FANTOM5^27^ enhancers, 331,601 super enhancers (SEdb^28^), 111,308 GWAS Catalog^31^ variants (with an association *P* ≤5.0×10^−8^; 9,229 with an independent replication), 2,191 GWAS variants fine-mapped across population biobanks with a posterior inclusion probability of causality≥0.9^32^, and 100,530 CNVs from a CNV morbidity map of developmental delay^50,51^.

To assess the correlation between constraint Z score and the collected functional elements, we intersected each annotation with the scored 1kb windows binned by constraint Z score (<-4, [−4,−3), [−3,−2), [−2,−1), [−1,−0), [0,1), [1,2), [2,3), [3,4), ≥4), and counted the frequency of overlapping windows within each bin. The enrichment of a given annotation (except CNVs) at a constraint level was evaluated by comparing the corresponding frequency to the genome-wide average using a Fisher’s exact test. In the analysis of CNVs, we assessed their enrichment in constrained regions by assigning each CNV the highest Z score among its overlapping windows and comparing the proportions of constrained CNVs (Z≥4) from cases of developmental delay and healthy controls. The enrichment was further examined using a logistic regression model to adjust for the size and gene content (gene constraint^1^ and gene number) of each CNV. We note that we performed all above analyses restricting to exclusively non-coding windows to evaluate the use of our constraint metric in characterizing the non-coding genome.

### Estimation of constraint for aggregated regulatory annotations

We estimated how constrained the sequences encoding regulatory elements overall compared to coding exons by aggregating the regulatory annotations at a 1kb scale. These included 7,246 promoter-, 154,003 enhancer-, 117 microRNA (miRNA)-, and 414,084 long non-coding RNA (lncRNA)-1kb elements, created from concatenating ENCODE cCREs-PLS, cCREs-dELS, GENCODE^73^ miRNA, and FANTOM6^74^ lncRNA annotations, respectively, into 1kb windows. Similarly, 27,875 exonic 1kb elements were created from aggregating all protein-coding exons. Constraint Z scores were computed for the created 1kb elements and the percentiles of each regulatory annotation were compared against the exonic region. Benchmarking on the 50^th^ percentile (median) of exonic regions, we estimated the proportion of the regulatory elements that are under selection as strong as the coding exons.

### Incorporation of constraint Z score into GWAS fine-mapping

To demonstrate the use of our constraint metric in statistical fine-mapping, we performed approximate functionally informed fine-mapping^39^ incorporating constraint Z score and our previous fine-mapping results for 119 UK Biobank (UKBB) traits^32^. The constraint Z scores were normalized and used as functional prior probabilities to update the posterior inclusion probabilities (PIPs; denoted as PIP_z_) based on the previous UKBB fine-mapping (using a uniform prior, PIP_unif_) and SuSiE^75^. To exclude signals that potentially correspond to coding variants, we restricted our analysis to 60,121 non-coding variants in 6,592 SuSiE 95% credible set (CS)-trait pairs that do not contain variants within 1 kb of exonic regions. A total of 13,069 variant-trait pairs were predicted to have an increased PIP (ΔPIP≥0.01) of causality. The variants, associated traits, and PIP scores (PIP_unif_ and PIP_z_) are provided in Supplementary Data 4.

### Comparison of constraint Z score and other predictive scores

We compared our constraint metric with other seven genome-wide predictive scores – Orion^13^, CDTS^14^, gwRVIS^18^, DR^15^, phyloP^20^, phastCons^19^, and GERP^41^. Each score was downloaded from the original study, lifted over to GRCh38 coordinates (for Orion) and multiplied by −1 (for CDTS, gwRVIS, and DR) when needed so that a higher value represents a higher constraint/conservation for all metrics. Pairwise correlation between the scores was assessed by comparing the mean value of each score on 1kb windows, using a Spearman’s rank correlation test.

We evaluated the predictive performance of each metric in distinguishing functional non-coding variants (“positive” variant set) from background variants (“negative” variant set). Four positive variant sets were compiled from public databases: 1) 9,229 variants from GWAS Catalog^31^ (with an independent replication), 2) 2,191 variants from a recent fine-mapping study^32^ (with a posterior inclusion probability of causality≥0.9), 3) 140 high-confidence variants from 2), and 4) 288 variants from ClinVar^40^ (annotated as “pathogenic”). All variants were filtered to non-coding regions; in particular, ClinVar variants were more strictly filtered to intergenic/intron variants given its strong predominance of variants close to proteincoding exons (>90% were splice site/region variants). A further stringent non-coding subset was generated by excluding variants within 10kb to any exons, which resulted in 1) 4,379, 2) 967, 3) 59, and 4) 7 variants. For each positive variant set, a negative variant set was created by randomly drawing variants from gnomAD (to ~10× the size of corresponding positive variant set), of which the most severe molecular consequence is intergenic or intron and the AF approximates the positive variant set; AF>5% and allele count (AC)=1 were applied respectively for matching positive variant set 1)-3) and 4), based on their AF distributions in gnomAD (Fig. 3b). The selected variants were scored by each of the eight metrics, using bedtools^72^ (for .bed files) and bigWigAverageOverBed^69^ (for .BigWig files), and the performance of each metric in classifying positive and negative variants was assessed by the area under curve (AUC) statistic, as presented by the receiver operating characteristic (ROC) curve.

To investigate whether different metrics capture complementary information in the classification, we fitted logistic regression models using all eight metrics as independent variables. The relative contribution of each metric was evaluated by the dominance analysis^76,77^, which estimates the dominance of one predictor over another by comparing their additional *R*^2^ contributions across all subset models. We further explored whether specific features were particularly captured by (and may have contributed to the performance of) our metric. We merged all positive variant sets and focused on a set of variants (N=204) that were uniquely prioritized by our metric, defined as being captured in the 99^th^ percentile of constraint Z score but not in that of any other scores. Specific features associated with these variants were evaluated by comparing values of the 13 genomic features of these variants to the rest of the positive variant set. The fold change was used to indicate the extent to which a feature is distinguished in variants captured by constraint Z score from others.

### Correlation of constraint between non-coding regulatory elements and protein-coding genes

To examine whether constraint of non-coding regulatory elements informs the constraint of their target genes, we compared constraint Z scores of enhancers linked to constrained genes and unconstrained genes. The former included well-established gene sets of 189 ClinGen^56^ haploinsufficient genes, 2,454 MGI^57^ essential genes mapped to human orthologs, 1, 771 OMIM^58^ autosomal dominant genes, and 1,920 LOEUF^1^ first-decile genes; and the latter included a curated list of 356 olfactory receptor genes and 189 LOEUF last-decile genes with at least 10 expected LoF variants (which are sufficiently powered to be classified into the most constrained decile^1^). The LOEUF underpowered list included 1,117 genes with ≤5 expected LoF variants. Enhancers linked to each gene were obtained from the Roadmap Epigenomics Enhancer-Gene Linking database, which used correlated patterns of activity between histone modifications and gene expression to predict enhancer-gene links^78,79^. For each gene, we aggregated and merged enhancers predicted from all 127 reference epigenomes and assigned the most constrained enhancer to each gene for the analysis of enhancer-gene constraint correlation (Supplementary Data 5).

In the analysis of correlation between tissue-specific enhancer constraint and tissue-specific gene expression, we processed the enhancer-gene links with the same principle as described above but within specific tissue types (as defined in the Roadmap Epigenomics metadata^55^). For each gene and tissue type, we searched for tissue-specific gene expression in the Genotype-Tissue Expression (GTEx^59^) database (RNASeQCv1.1.9) and computed a normalized median expression for each gene (log2(TPM+1)). Enhancer constraint and gene expression values were calculated for 11 matched tissue types, and the correlation within each tissue type was evaluated by regressing gene expression on enhancer constraint, including gene constraint (LOEUF score) as a covariate.

### Incorporation of non-coding constraint of regulatory elements into gene constraint modeling

To demonstrate the practical value of non-coding constraint in improving gene constraint modeling, we compared two models – using 1) LOEUF and 2) LOEUF+enhancer constraint Z score (as described in Correlation of constraint between non-coding regulatory elements and protein-coding genes) – in predicting constrained genes, with a particular focus on genes that were underpowered in LOEUF. A set of 3,220 unique constrained genes were curated from ClinGen^56^, MGI^57^, and OMIM^58^ (see Correlation of constraint between non-coding regulatory elements and protein-coding genes), and a set of 356 olfactory receptor genes was used as the unconstrained genes. We trained logistic regression models on 50% of the genes and tested the performance on 77 underpowered genes in the remaining 50%. The predictive performance of the two models were measured by AUC, and the significance of the difference in AUCs was assessed using a bootstrap test^80^.

### Power of constraint detection

We estimated the power of our metric in detecting non-coding constraint as the percentage of the non-coding genome to obtain a high constraint Z score (Z≥4) under a certain strength of negative selection, which was quantified by the level of depletion of variation (i.e., 1-observed/expected). For a given depletion of variation, the minimum number of expected variants to achieve a Z≥4 was determined, and the number of samples required to achieve the expected number of variants was estimated using a linear model of log(number of expected variants) ~ log(number of samples) from downsampling the gnomAD dataset. The power was estimated at two scales – 1kb (used in this study) and 100bp – and benchmarked by the depletion of variation observed in coding exons of similar size.

---

**Extended Data Fig. 1:**
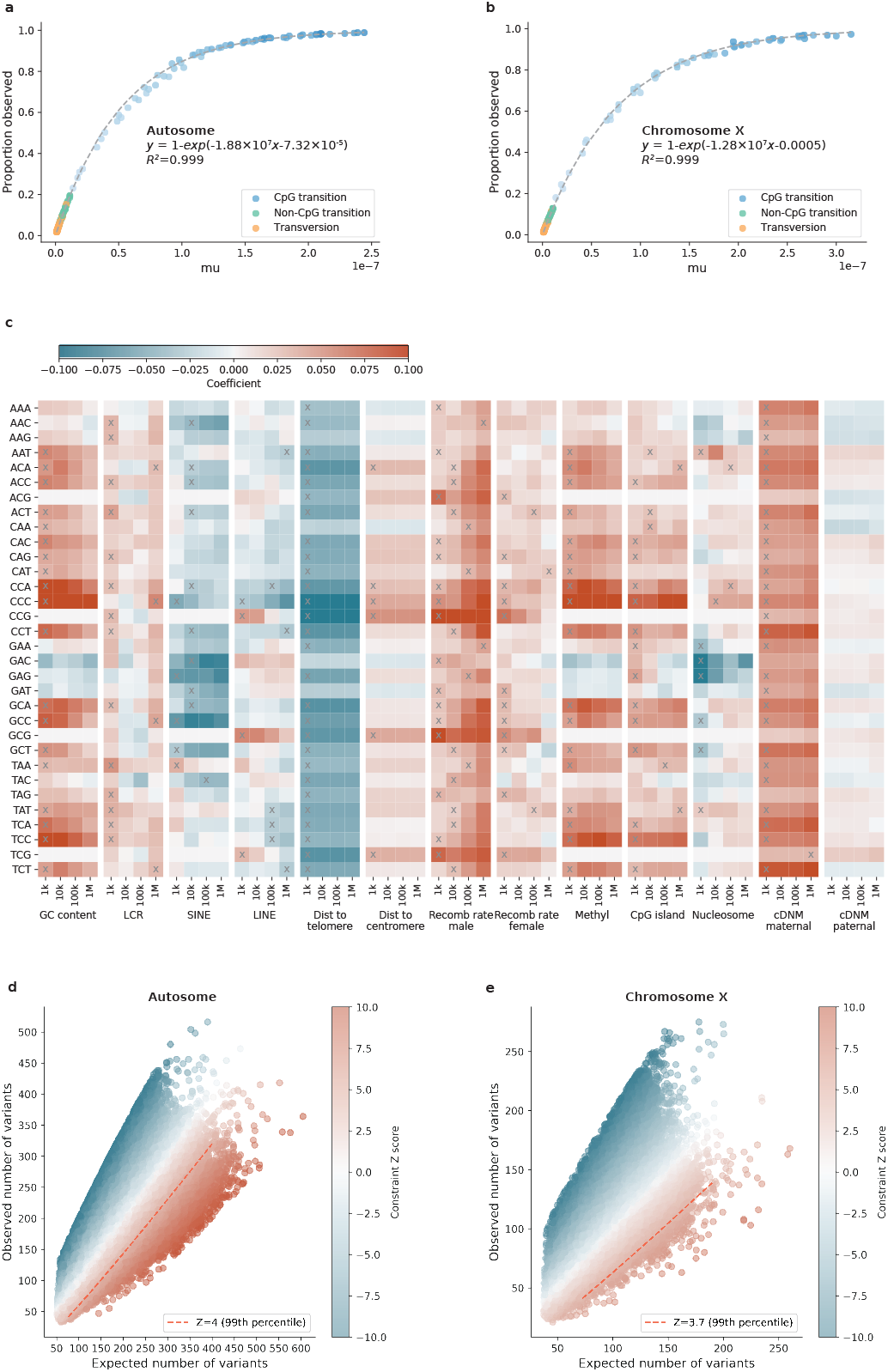
Construction of mutational model and constraint Z score. **a**,**b**, Estimation of trinucleotide context-specific mutation rates. The proportion of possible variants observed for each substitution and context in 76,156 gnomAD genomes (y-axis) is exponentially correlated with the absolute mutation rate estimated from 1,000 downsampled genomes (x-axis). Fit lines were modeled separately for human autosomes (**a**) and chromosome X (**b**). **c**, Estimation of the effects of regional genomic features on mutation rates. The effects of 13 genomic features at four scales (window sizes 1kb-1Mb; x-axis) on the mutation rate of 32 trinucleotide contexts (y-axis) are shown, colored by the coefficient from regressing *de novo* mutations (DNMs) on each specific feature and window size. Red/Blue color indicates a positive/negative effect of increasing the feature value on mutation rates; grey crosses indicate significant features at the smallest possible window size after Bonferroni correction for 13×4=52 tests. Abbreviations: LCR=low-complexity region, SINE/LINE=short/long interspersed nuclear element, Dist=Distance, Recomb=Recombination, Methyl=Methylation. **d**,**e**, The distribution of constraint Z score as a function of expected and observed variation. Each point represents the Z score of a 1kb window on the genome (N=1,984,900 on autosomes (**d**) and N=57,729 on chromosome X (**e**)), which quantifies the deviation of observed variation from expectation. A positive Z score (red) indicates depletion of variation (observed<expected) and the higher the Z score the stronger the depletion; the red dashed line indicates the 99^th^ percentile of Z scores across the autosomes (**d**) or chromosome X (**e**).

**Extended Data Fig. 2:**
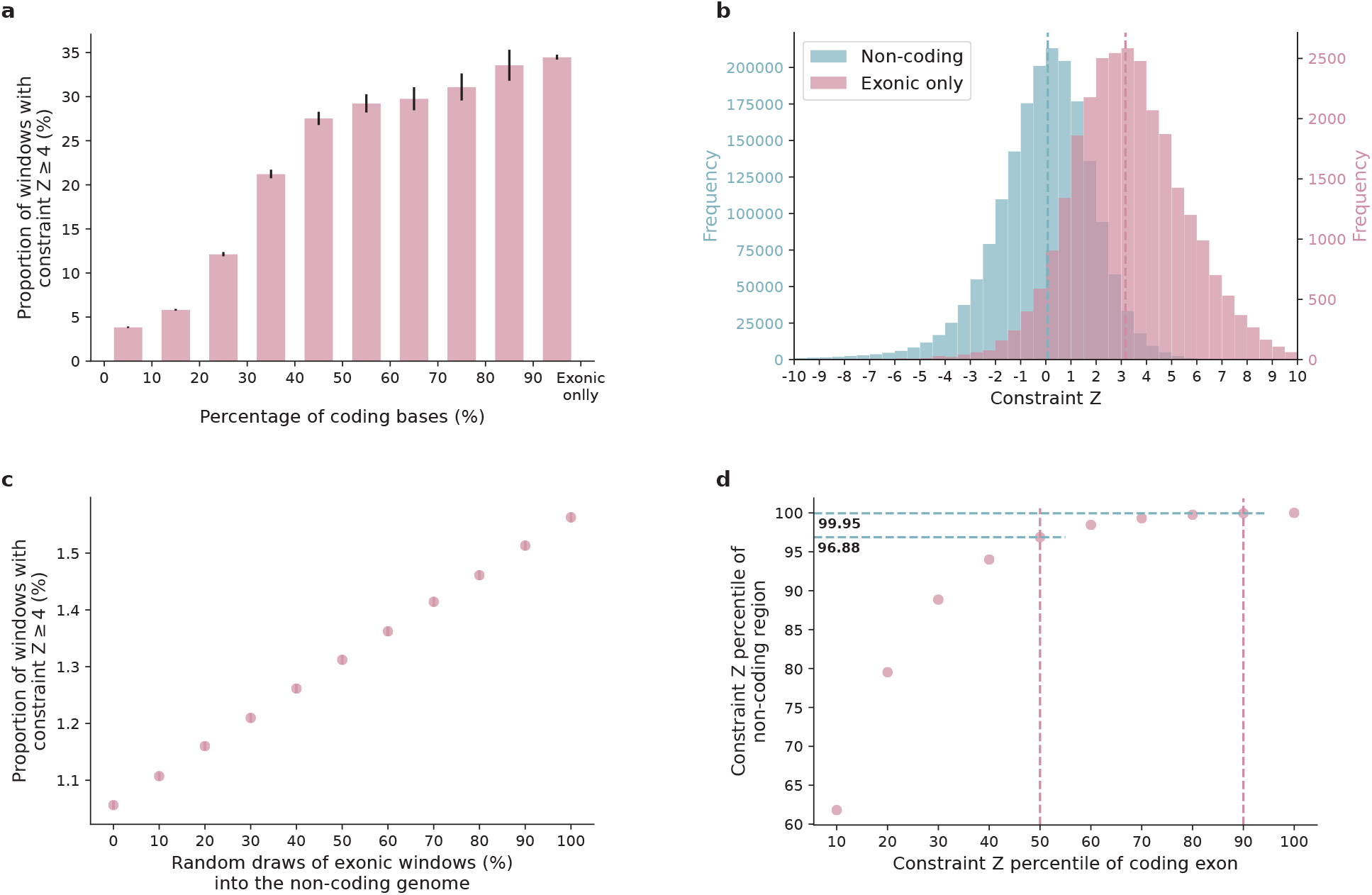
Comparison of constraint Z score between coding and non-coding regions. **a**, The proportion of highly constrained windows (Z≥4) as a function of the percentage of coding sequences in a window. The intervals (x-axis) are left exclusive and right inclusive. “Exonic only” refers to the 1kb windows created from directly concatenating coding exons into 1kb sequences. **b**, The exonic-only regions (N=27,875; purple) present a significantly higher constraint Z score than regions that are exclusively noncoding (N=1,843,559; blue). Dashed lines indicate the medians. **c**, The proportion of highly constrained windows (Z≥4) as a function of the proportion of exonic windows being added to the dataset of noncoding windows. **d**, Constraint Z score percentiles of non-coding versus exonic windows. About 0.05% (100-99.95%) and 3.12% (100-96.88%) of the non-coding windows exhibit similar constraint to the 90^th^ and 50^th^ of exonic regions, respectively.

**Extended Data Fig. 3:**
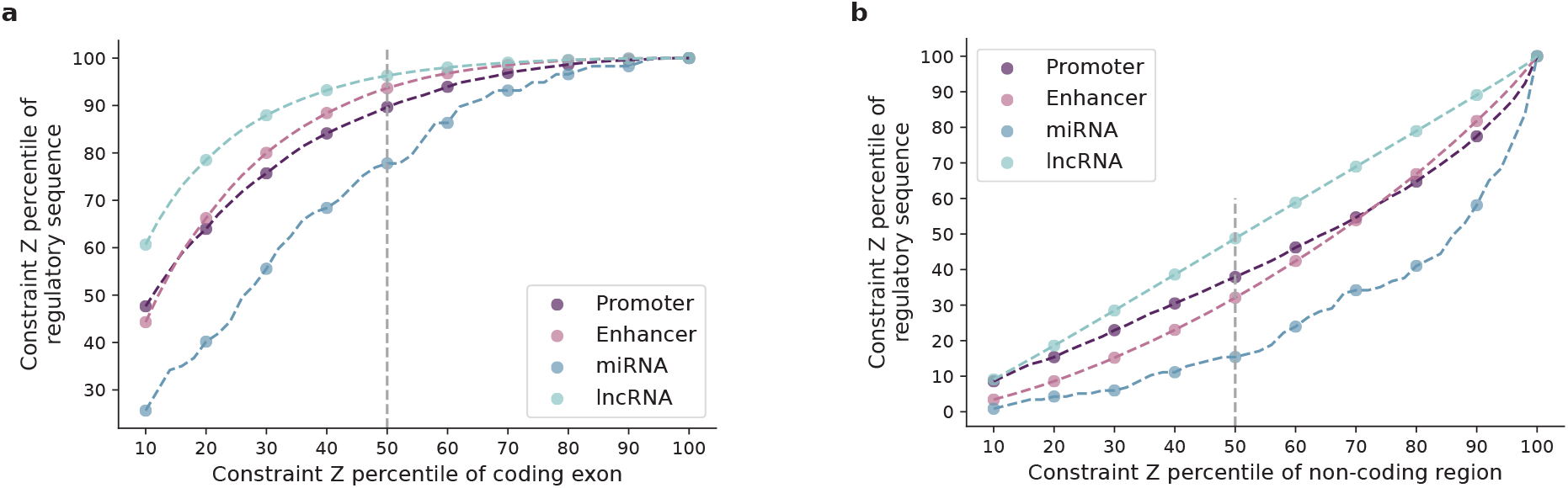
Estimation of constraint for aggregated regulatory annotations. **a**,**b**, Constraint Z scores of aggregated promoter (dark purple), enhancer (light purple), microRNA (miRNA; dark blue), and long non-coding RNA (lncRNA; light blue) annotations are compared against those of exonic (**a**) and noncoding (**b**) regions at a 1kb scale. The constraint Z score percentiles of each annotation (y-axis) are benchmarked by the score deciles of exonic or non-coding regions (10-100 percentiles; x-axis); the grey dashed vertical line indicates the median (50^th^ percentile).

**Extended Data Fig. 4:**
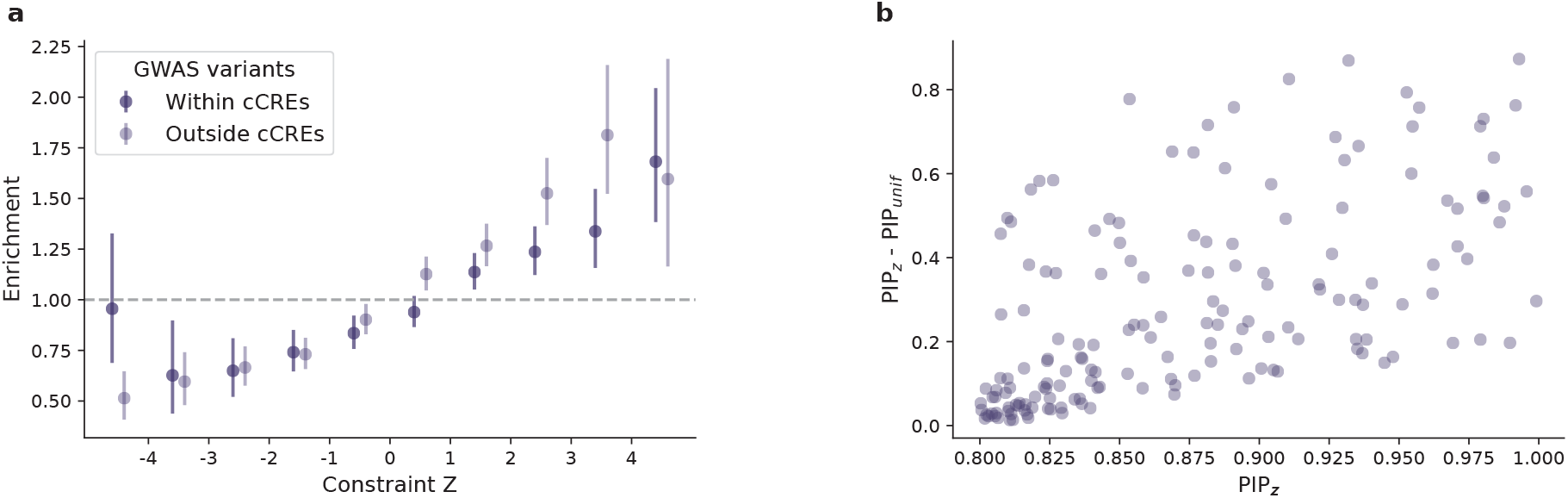
Applications of constraint Z score for characterizing non-coding regions in addition to existing functional annotations. **a**, Use of constraint Z score for prioritizing non-coding regions with or without a regulatory annotation. Constrained non-coding regions are enriched for GWAS variants, independent of the candidate cis-regulatory element (cCRE) annotation from ENCODE. **b**, Use of constraint Z score in statistical fine-mapping. The increase in posterior inclusion probability (PIP) when incorporating constraint Z score as a functional prior into previous fine-mapping results (that used a uniform prior; denoted as PIP_z_ and PIP_unif_, respectively) is shown for 164 new likely causal associations with a PIP_z_ ≥0.8 as a function of PIP_z_.

**Extended Data Fig. 5:**
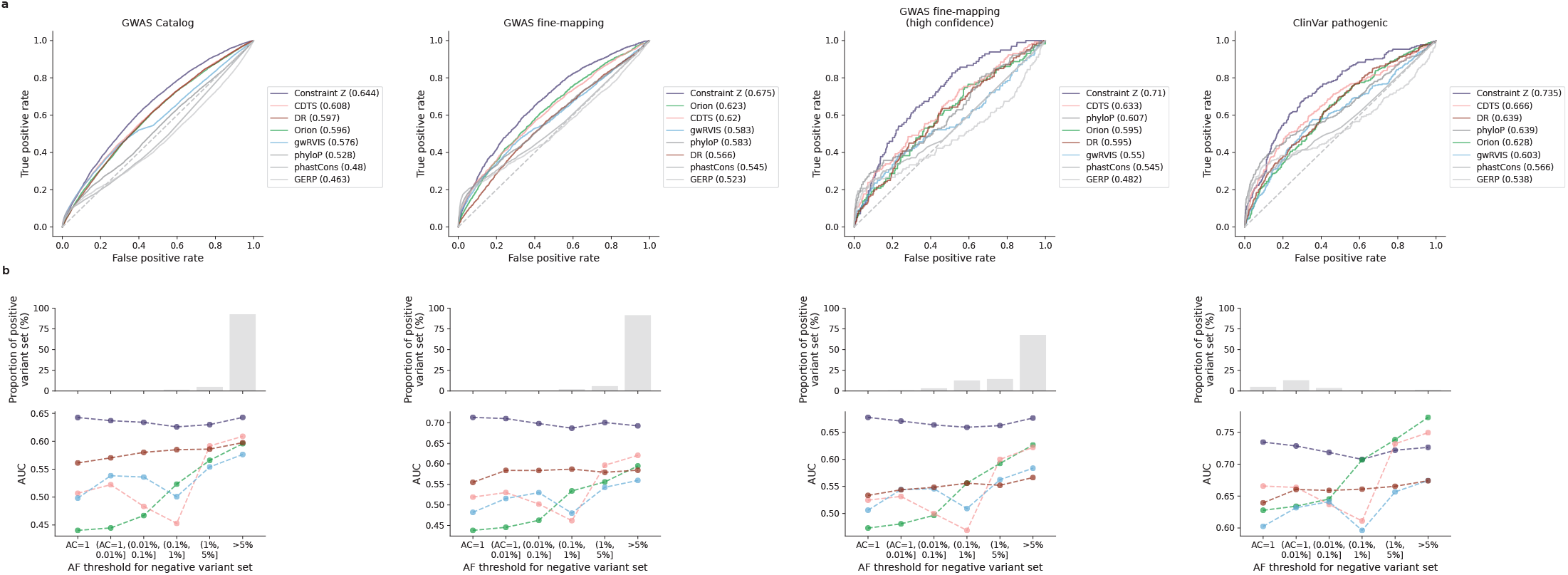
Comparison of constraint Z score and other predictive scores in prioritizing noncoding variants. **a**, Receiver operating characteristic (ROC) curves of constraint Z score and other seven metrics in classifying putative functional non-coding variants (“positive” variant set) – left to right: 9,229 GWAS Catalog variants, 2,191 GWAS fine-mapping variants, a subset of 140 high-confidence fine-mapped variants, and 288 ClinVar pathogenic variants – against “negative” variant set randomly drew from the population with a similar allele frequency (AF). AF>5% and allele count (AC)=1 were applied respectively for matching the three GWAS variant sets and the ClinVar variant set, based on their AF distributions in gnomAD (shown in **b**). **b**, AUCs of the classification with a varying AF threshold for the negative variant set. As most GWAS variants are common and most ClinVar variants are very rare (not seen in the population), AF>5% and AC=1 were applied respectively in the primary analyses shown in **a**.

**Extended Data Fig. 6:**
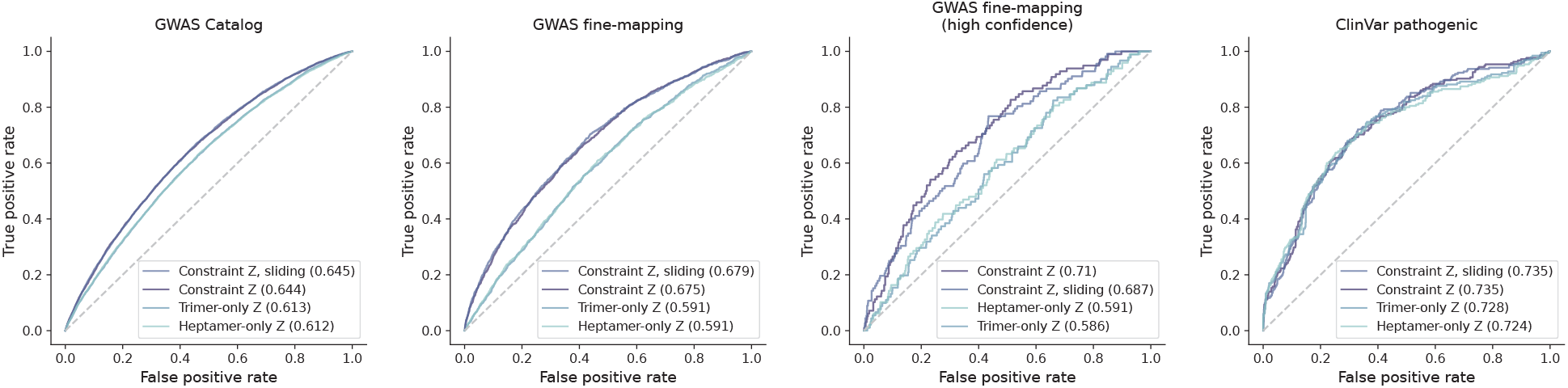
Comparison of constraint Z scores built from different mutational models and genomic windows. Our constraint Z score (presented in this study) outperforms the scores rebuilt from mutational models that only consider local sequence context – trinucleotide (trimer-only) or heptanucleotide (heptamer-only) – without adjustment on mutation rate by regional genomic features, and the performance is robust to the artificial break of genomic windows when computed at a 1kb sliding by 100bp scale.

**Extended Data Fig. 7:**
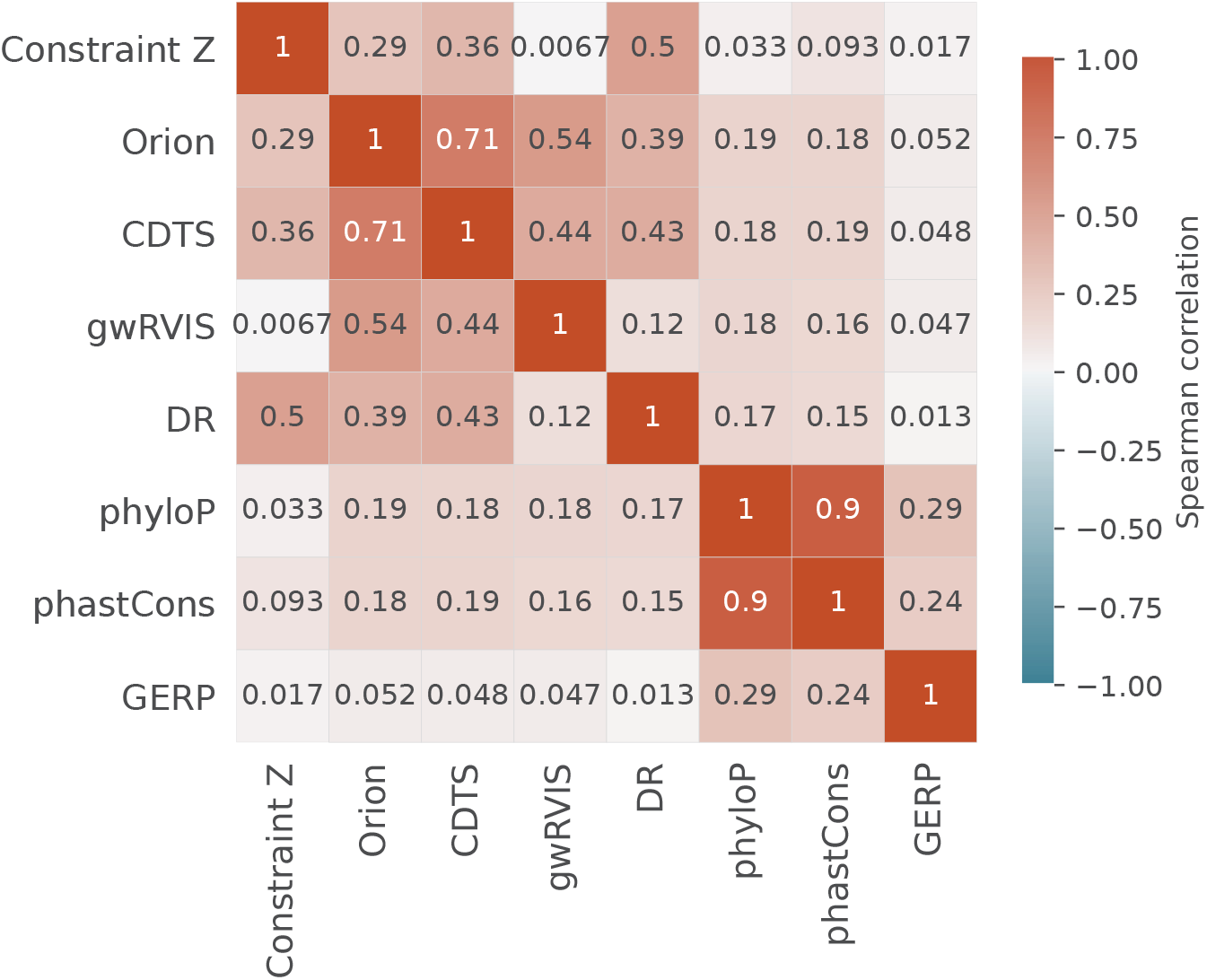
Pairwise correlations between different constraint/conservation metrics. The Spearman’s rank correlation between each pair of the eight metrics was computed based on the mean value of each score on 1kb windows across the genome.

**Extended Data Fig. 8:**
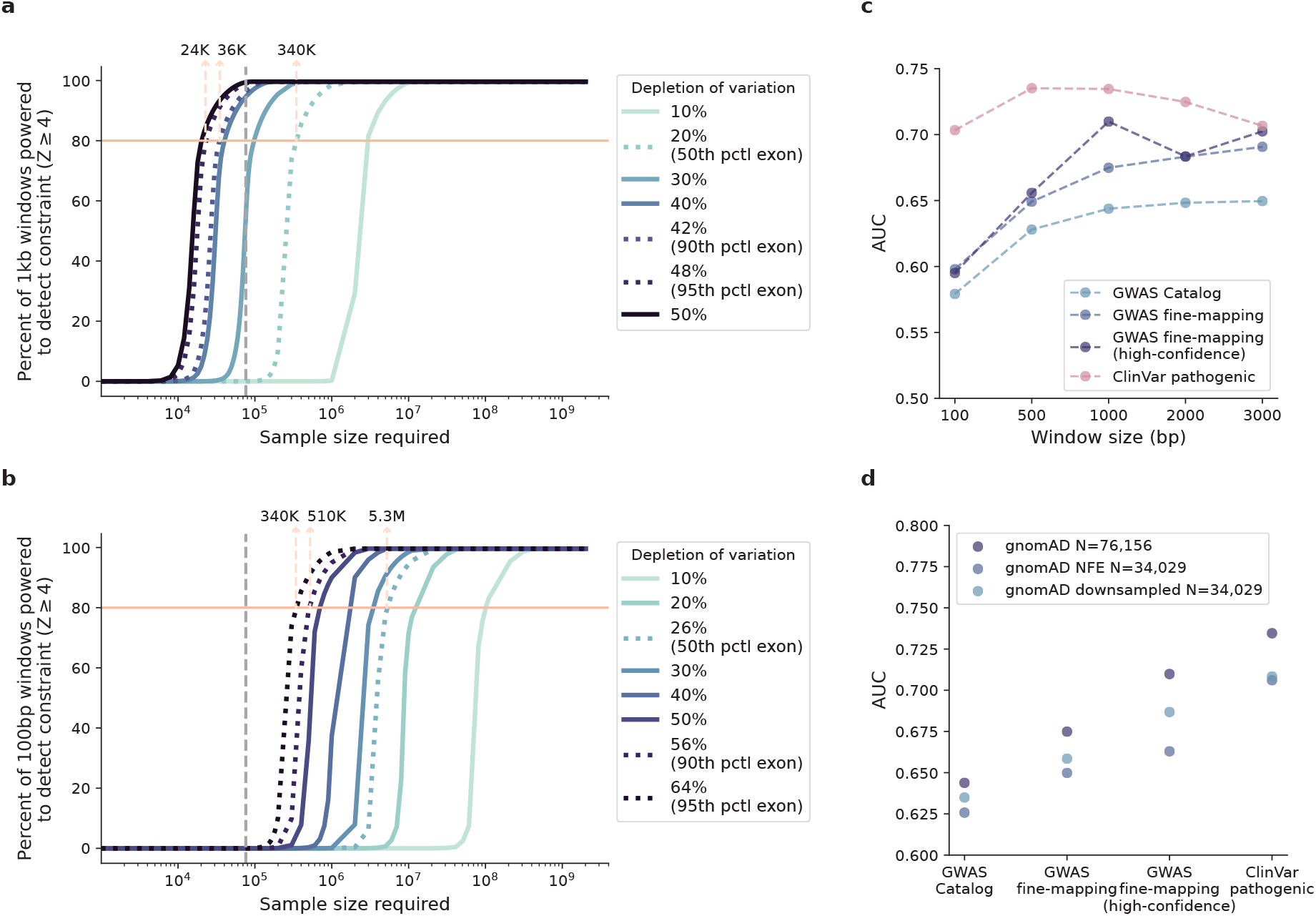
Power of constraint detection. **a**,**b**, The sample size required for well-powered noncoding constraint detection. The percentage of non-coding regions powered to detect constraint (Z≥4) at a 1kb (**a**) and 100bp (**b**) scale under varying levels of selection (depletion of variation) is shown as a function of log-scaled sample size. Lighter color indicates milder deletion of variation (weaker selection), which requires a larger sample size to detect constraint; the grey dashed vertical line indicates the current sample size of 76,156 genomes. Dotted curves (left to right) benchmark the 95^th^, 90^th^, and 50^th^ percentile of depletion of variation observed in coding exons of similar size. The number of samples required to obtain an 80% detection power is labeled at corresponding benchmarks. **c**, AUCs of constraint Z scores computed on different window sizes in identifying putative functional non-coding variants. 1kb (used in this study) presents the optimal window size with high performance while maintaining reasonable resolution. **d**, AUCs of constraint Z scores computed from different subsets of gnomAD in identifying putative functional non-coding variants. While with an equal sample size, the downsampled dataset with diverse ancestries presents higher performance than the Non-Finnish European (NFE)-only dataset.

## Genome Aggregation Database Consortium

Maria Abreu^1^, Carlos A. Aguilar Salinas^2^, Tariq Ahmad^3^, Christine M. Albert^4,5^, Jessica Alföldi^6,7^, Diego Ardissino^8^, Irina M. Armean^6,7,9^, Gil Atzmon^10,11^, Eric Banks^12^, John Barnard^13^, Samantha M. Baxter^6^, Laurent Beaugerie^14^, Emelia J. Benjamin^15,16,17^, David Benjamin^12^, Louis Bergelson^12^, Michael Boehnke^18^, Lori L. Bonnycastle^19^, Erwin P. Bottinger^20^, Donald W. Bowden^21^,^22^,^23^, Matthew J. Bown^24^,^25^, Steven Brant^26^, Sarah E. Calvo^6^,^27^, Hannia Campos^28^,^29^, John C. Chambers^30^,^31^,^32^, Juliana C.N. Chan^33^,^109^,^110^, Katherine R. Chao^6^, Sinéad Chapman^6^,^7^,^34^, Daniel Chasman^4,35^, Siwei Chen^6,7^, Rex L. Chisholm^36^, Judy Cho^20^, Rajiv Chowdhury^37^, Mina K. Chung^38^, Wendy K. Chung^39,40,41^, Kristian Cibulskis^12^, Bruce Cohen^35,42^, Ryan L. Collins^6,27,43^, Kristen M. Connolly^44^, Adolfo Correa^45^, Miguel Covarrubias^12^, Beryl Cummings^6,43^, Dana Dabelea^46^, Mark J. Daly^6,7,47^, John Danesh^37^, Dawood Darbar^48^, Joshua Denny^49^, Stacey Donnelly^6^, Ravindranath Duggirala^50^, Josée Dupuis^51,52^, Patrick T. Ellinor^6,53^, Roberto Elosua^54,55,56^, James Emery^12^, Eleina England^6^, Jeanette Erdmann^57,58,59^, Tõnu Esko^6,60^, Emily Evangelista^6^, Yossi Farjoun^12^, Diane Fatkin^61,62,63^, Steven Ferriera^64^, Jose Florez^35,65,66^, Laurent C. Francioli^6,7^, Andre Franke^67^, Martti Färkkilä^68^, Stacey Gabriel^64^, Kiran Garimella^12^, Laura D. Gauthier^12^, Jeff Gentry^12^, Gad Getz^35^,^69^,^70^, David C. Glahn^71,72^, Benjamin Glaser^73^, Stephen J. Glatt^74^, David Goldstein^75,76^, Clicerio Gonzalez^77^, Julia K. Goodrich^6^, Leif Groop^78,79^, Sanna Gudmundsson^6,7,80^, Namrata Gupta^6,64^, Andrea Haessly^12^, Christopher Haiman^205^, Ira Hall^82^, Craig Hanis^83^, Matthew Harms^84^,^85^, Mikko Hiltunen^86^, Matti M. Holi^87^, Christina M. Hultman^88^,^89^, Chaim Jalas^90^, Thibault Jeandet^12^, Mikko Kallela^91^, Diane Kaplan^12^, Jaakko Kaprio^79^, Konrad J. Karczewski^6,7,34^, Sekar Kathiresan^27^,^35^,^92^, Eimear Kenny^89^,^93^, Bong-Jo Kim^94^, Young Jin Kim^94^, George Kirov^95^, Zan Koenig^6^, Jaspal Kooner^31^,^96^,^97^, Seppo Koskinen^98^, Harlan M. Krumholz^99^, Subra Kugathasan^100^, Soo Heon Kwak^101^, Markku Laakso^102,103^, Nicole Lake^104^, Trevyn Langsford^12^, Kristen M. Laricchia^6,7^, Terho Lehtimäki^105^, Monkol Lek^104^, Emily Lipscomb^6^, Christopher Llanwarne^12^, Ruth J.F. Loos^20,106^, Steven A. Lubitz^6,53^, Teresa Tusie Luna^107,108^, Ronald C.W. Ma^33^,^109^,^110^, Daniel G. MacArthur^6^,^111^,^112^, Gregory M. Marcus^113^, Jaume Marrugat^55^,^114^, Alicia R. Martin^6^, Kari M. Mattila^105^, Steven McCarroll^34^,^115^, Mark I. McCarthy^116^,^117^,^118^, Jacob McCauley^119^,^120^, Dermot McGovern^121^, Ruth McPherson^122^, James B. Meigs^6,35,123^, Olle Melander^124^, Andres Metspalu^125^, Deborah Meyers^126^, Eric V. Minikel^6^, Braxton D, Mitchell^127^, Vamsi K. Mootha^6,128^, Ruchi Munshi^12^, Aliya Naheed^129^, Saman Nazarian^130,131^, Benjamin M. Neale^6,7^, Peter M. Nilsson^132^, Sam Novod^12^, Anne H. O’Donnell-Luria^6,7,80^, Michael C. O’Donovan^95^, Yukinori Okada^133^,^134^,^135^, Dost Ongur^35^,^42^, Lorena Orozco^136^, Michael J. Owen^95^, Colin Palmer^137^, Nicholette D. Palmer^138^, Aarno Palotie^7^,^34^,^79^, Kyong Soo Park^101^,^139^, Carlos Pato^140^, Nikelle Petrillo^12^, William Phu^6^,^80^, Timothy Poterba^6^,^7^,^34^, Ann E. Pulver^141^, Dan Rader^130^,^142^, Nazneen Rahman^143^, Heidi L. Rehm^6^,^27^, Alex Reiner^144^,^145^, Anne M. Remes^146^, Dan Rhodes^6^, Stephen S. Rich^147,148^, John D. Rioux^149,150^, Samuli Ripatti^79,151,152^, David Roazen^12^, Dan M. Roden^153^,^154^, Jerome I. Rotter^155^, Valentin Ruano-Rubio^12^, Nareh Sahakian^12^, Danish Saleheen^156^,^157^,^158^, Veikko Salomaa^159^, Andrea Saltzman^6^, Nilesh J. Samani^24,25^, Kaitlin E. Samocha^6,27^, Jeremiah Scharf^6,27,34^, Molly Schleicher^6^, Heribert Schunkert^160^,^161^, Sebastian Schönherr^162^, Eleanor Seaby^6^, Cotton Seed^7^,^34^, Svati H. Shah^163^, Megan Shand^12^, Moore B. Shoemaker^164^, Tai Shyong^165,166^, Edwin K. Silverman^167,168^, Moriel Singer-Berk^6^, Pamela Sklar^169^,^170^,^171^, J. Gustav Smith^152^,^172^,^173^, Jonathan T. Smith^12^, Hilkka Soininen^174^, Harry Sokol^175^,^176^,^177^, Matthew Solomonson^6,7^, Rachel G. Son^6^, Jose Soto^12^, Tim Spector^178^, Christine Stevens^6,7,34^, Nathan Stitziel^82,179^, Patrick F. Sullivan^88^,^180^, Jaana Suvisaari^159^, E. Shyong Tai^181^,^182^,^183^, Michael E. Talkowski^6^,^27^,^34^, Yekaterina Tarasova^6^, Kent D. Taylor^155^, Yik Ying Teo^181^,^184^,^185^, Grace Tiao^6^,^7^, Kathleen Tibbetts^12^, Charlotte Tolonen^12^, Ming Tsuang^186^,^187^, Tiinamaija Tuomi^79,188,189^, Dan Turner^190^, Teresa Tusie-Luna^191,192^, Erkki Vartiainen^193^, Marquis P. Vawter^194^, Christopher Vittal^6^,^7^, Gordon Wade^12^, Arcturus Wang^6^,^7^,^34^, Qingbo Wang^6^,^133^, James S. Ware^6^,^195^,^196^, Hugh Watkins^197^, Nicholas A. Watts^6,7^, Rinse K. Weersma^198^, Ben Weisburd^12^, Maija Wessman^79,199^, Nicola Whiffin^6^,^200^,^201^, Michael W. Wilson^6^,^7^, James G. Wilson^202^, Ramnik J. Xavier^203^,^204^, Mary T. Yohannes^6^

^1^University of Miami Miller School of Medicine, Gastroenterology, Miami, USA

^2^Unidad de Investigacion de Enfermedades Metabolicas, Instituto Nacional de Ciencias Medicas y Nutricion, Mexico City, Mexico

^3^Peninsula College of Medicine and Dentistry, Exeter, UK

^4^Division of Preventive Medicine, Brigham and Women’s Hospital, Boston, MA, USA

^5^Division of Cardiovascular Medicine, Brigham and Women’s Hospital and Harvard Medical School, Boston, MA, USA

^6^Program in Medical and Population Genetics, Broad Institute of MIT and Harvard, Cambridge, MA, USA

^7^Analytic and Translational Genetics Unit, Massachusetts General Hospital, Boston, MA, USA

^8^Department of Cardiology University Hospital, Parma, Italy

^9^European Molecular Biology Laboratory, European Bioinformatics Institute, Wellcome Genome Campus, Hinxton, Cambridge, UK

^10^Department of Biology Faculty of Natural Sciences, University of Haifa, Haifa, Israel

^11^Departments of Medicine and Genetics, Albert Einstein College of Medicine, Bronx, NY, USA

^12^Data Science Platform, Broad Institute of MIT and Harvard, Cambridge, MA, USA

^13^Department of Quantitative Health Sciences, Lerner Research Institute Cleveland Clinic, Cleveland, OH, USA ^14^Sorbonne Université, APHP, Gastroenterology Department Saint Antoine Hospital, Paris, France

^15^NHLBI and Boston University’s Framingham Heart Study, Framingham, MA, USA

^16^Department of Medicine, Boston University School of Medicine, Boston, MA, USA

^17^Department of Epidemiology, Boston University School of Public Health, Boston, MA, USA

^18^Department of Biostatistics and Center for Statistical Genetics, University of Michigan, Ann Arbor, MI, USA

^19^National Human Genome Research Institute, National Institutes of Health Bethesda, MD, USA

^20^The Charles Bronfman Institute for Personalized Medicine, Icahn School of Medicine at Mount Sinai, New York, NY, USA

^21^Department of Biochemistry, Wake Forest School of Medicine, Winston-Salem, NC, USA

^22^Center for Genomics and Personalized Medicine Research, Wake Forest School of Medicine, Winston-Salem, NC, USA

^23^Center for Diabetes Research, Wake Forest School of Medicine, Winston-Salem, NC, USA

^24^Department of Cardiovascular Sciences, University of Leicester, Leicester, UK

^25^NIHR Leicester Biomedical Research Centre, Glenfield Hospital, Leicester, UK

^26^John Hopkins Bloomberg School of Public Health, Baltimore, MD, USA

^27^Center for Genomic Medicine, Massachusetts General Hospital, Boston, MA, USA

^28^Harvard School of Public Health, Boston, MA, USA

^29^Central American Population Center, San Pedro, Costa Rica

^30^Department of Epidemiology and Biostatistics, Imperial College London, London, UK

^31^Department of Cardiology, Ealing Hospital, NHS Trust, Southall, UK

^32^Imperial College, Healthcare NHS Trust Imperial College London, London, UK

^33^Department of Medicine and Therapeutics, The Chinese University of Hong Kong, Hong Kong, China

^34^Stanley Center for Psychiatric Research, Broad Institute of MIT and Harvard, Cambridge, MA, USA

^35^Department of Medicine, Harvard Medical School, Boston, MA, USA

^36^Northwestern University Feinberg School of Medicine, Chicago, IL, USA

^37^University of Cambridge, Cambridge, England

^38^Departments of Cardiovascular, Medicine Cellular and Molecular Medicine Molecular Cardiology, Quantitative Health Sciences, Cleveland Clinic, Cleveland, OH, USA

^39^Department of Pediatrics, Columbia University Irving Medical Center, New York, NY, USA

^40^Herbert Irving Comprehensive Cancer Center, Columbia University Medical Center, New York, NY, USA

^41^Department of Medicine, Columbia University Medical Center, New York, NY, USA

^42^McLean Hospital, Belmont, MA, USA

^43^Division of Medical Sciences, Harvard Medical School, Boston, MA, USA

^44^Genomics Platform, Broad Institute of MIT and Harvard, Cambridge, MA, USA

^45^Department of Medicine, University of Mississippi Medical Center, Jackson, MI, USA

^46^Department of Epidemiology Colorado School of Public Health Aurora, CO, USA

^47^Institute for Molecular Medicine Finland, (FIMM) Helsinki, Finland

^48^Department of Medicine and Pharmacology, University of Illinois at Chicago, Chicago, IL, USA

^49^Vanderbilt University Medical Center, Nashville, TN, USA

^50^Department of Genetics, Texas Biomedical Research Institute, San Antonio, TX, USA

^51^Department of Biostatistics, Boston University School of Public Health, Boston, MA, USA

^52^National Heart Lung and Blood Institute’s Framingham Heart Study, Framingham, MA, USA

^53^Cardiac Arrhythmia Service and Cardiovascular Research Center, Massachusetts General Hospital, Boston, MA, USA

^54^Cardiovascular Epidemiology and Genetics Hospital del Mar Medical Research Institute, (IMIM) Barcelona Catalonia, Spain

^55^CIBER CV Barcelona, Catalonia, Spain

^56^Departament of Medicine, Medical School University of Vic-Central, University of Catalonia, Vic Catalonia, Spain

^57^Institute for Cardiogenetics, University of Lübeck, Lübeck, Germany

^58^German Research Centre for Cardiovascular Research, Hamburg/Lübeck/Kiel, Lübeck, Germany

^59^University Heart Center Lübeck, Lübeck, Germany

^60^Estonian Genome Center, Institute of Genomics University of Tartu, Tartu, Estonia

^61^Victor Chang Cardiac Research Institute, Darlinghurst, NSW, Australia

^62^Faculty of Medicine, UNSW Sydney, Kensington, NSW, Australia

^63^Cardiology Department, St Vincent’s Hospital, Darlinghurst, NSW, Australia

^64^Broad Genomics, Broad Institute of MIT and Harvard, Cambridge, MA, USA

^65^Diabetes Unit and Center for Genomic Medicine, Massachusetts General Hospital, Boston, MA, USA

^66^Programs in Metabolism and Medical & Population Genetics, Broad Institute of MIT and Harvard, Cambridge, MA, USA

^67^Institute of Clinical Molecular Biology, (IKMB) Christian-Albrechts-University of Kiel, Kiel, Germany

^68^Helsinki University and Helsinki University Hospital Clinic of Gastroenterology, Helsinki, Finland

^69^Bioinformatics Program MGH Cancer Center and Department of Pathology, Boston, MA, USA

^70^Cancer Genome Computational Analysis, Broad Institute of MIT and Harvard, Cambridge, MA, USA

^71^Department of Psychiatry and Behavioral Sciences, Boston Children’s Hospitaland Harvard Medical School, Boston, MA, USA

^72^Harvard Medical School Teaching Hospital, Boston, MA, USA

^73^Department of Endocrinology and Metabolism, Hadassah Medical Center and Faculty of Medicine, Hebrew University of Jerusalem, Israel

^74^Department of Psychiatry and Behavioral Sciences, SUNY Upstate Medical University, Syracuse, NY, USA

^75^Institute for Genomic Medicine, Columbia University Medical Center Hammer Health Sciences, New York, NY, USA

^76^Department of Genetics & Development Columbia University Medical Center, Hammer Health Sciences, New York, NY, USA

^77^Centro de Investigacion en Salud Poblacional, Instituto Nacional de Salud Publica, Mexico

^78^Lund University Sweden, Sweden

^79^Institute for Molecular Medicine Finland, (FIMM) HiLIFE University of Helsinki, Helsinki, Finland

^80^Division of Genetics and Genomics, Boston Children’s Hospital, Boston, MA, USA

^81^Lund University Diabetes Centre, Malmö, Skåne County, Sweden

^82^Washington School of Medicine, St Louis, MI, USA

^83^Human Genetics Center University of Texas Health Science Center at Houston, Houston, TX, USA

^84^Department of Neurology Columbia University, New York City, NY, USA

^85^Institute of Genomic Medicine, Columbia University, New York City, NY, USA

^86^Institute of Biomedicine, University of Eastern Finland, Kuopio, Finland

^87^Department of Psychiatry, Helsinki University Central Hospital Lapinlahdentie, Helsinki, Finland

^88^Department of Medical Epidemiology and Biostatistics, Karolinska Institutet, Stockholm, Sweden

^89^Icahn School of Medicine at Mount Sinai, New York, NY, USA

^90^Bonei Olam, Center for Rare Jewish Genetic Diseases, Brooklyn, NY, USA

^91^Department of Neurology, Helsinki University, Central Hospital, Helsinki, Finland

^92^Cardiovascular Disease Initiative and Program in Medical and Population Genetics, Broad Institute of MIT and Harvard, Cambridge, MA, USA

^93^Charles Bronfman Institute for Personalized Medicine, New York, NY, USA

^94^Division of Genome Science, Department of Precision Medicine, National Institute of Health, Republic of Korea

^95^MRC Centre for Neuropsychiatric Genetics & Genomics, Cardiff University School of Medicine, Cardiff, Wales

^96^Imperial College, Healthcare NHS Trust, London, UK

^97^National Heart and Lung Institute Cardiovascular Sciences, Hammersmith Campus, Imperial College London, London, UK

^98^Department of Health THL-National Institute for Health and Welfare, Helsinki, Finland

^99^Section of Cardiovascular Medicine, Department of Internal Medicine, Yale School of Medicine, Center for Outcomes Research and Evaluation Yale-New Haven Hospital, New Haven, CT, USA

^100^Division of Pediatric Gastroenterology, Emory University School of Medicine, Atlanta, GA, USA

^101^Department of Internal Medicine, Seoul National University Hospital, Seoul, Republic of Korea

^102^The University of Eastern Finland, Institute of Clinical Medicine, Kuopio, Finland

^103^Kuopio University Hospital, Kuopio, Finland

^104^Department of Genetics, Yale School of Medicine, New Haven, CT, USA

^105^Department of Clinical Chemistry Fimlab Laboratories and Finnish Cardiovascular Research Center-Tampere Faculty of Medicine and Health Technology, Tampere University, Finland

^106^The Mindich Child Health and Development, Institute Icahn School of Medicine at Mount Sinai, New York, NY, USA

^107^National Autonomous University of Mexico, Mexico City, Mexico

^108^Salvador Zubirán National Institute of Health Sciences and Nutrition, Mexico City, Mexico

^109^Li Ka Shing Institute of Health Sciences, The Chinese University of Hong Kong, Hong Kong, China

^110^Hong Kong Institute of Diabetes and Obesity, The Chinese University of Hong Kong, Hong Kong, China

^111^Centre for Population Genomics, Garvan Institute of Medical Research and UNSW Sydney, Sydney, Australia

^112^Centre for Population Genomics, Murdoch Children’s Research Institute, Melbourne, Australia

^113^University of California San Francisco Parnassus Campus, San Francisco, CA, USA

^114^Cardiovascular Research REGICOR Group, Hospital del Mar Medical Research Institute, (IMIM) Barcelona, Catalonia, Spain

^115^Department of Genetics, Harvard Medical School, Boston, MA, USA

^116^Oxford Centre for Diabetes, Endocrinology and Metabolism, University of Oxford, Churchill Hospital Old Road Headington, Oxford, OX, LJ, UK

^117^Welcome Centre for Human Genetics, University of Oxford, Oxford, OX, BN, UK

^118^Oxford NIHR Biomedical Research Centre, Oxford University Hospitals, NHS Foundation Trust, John Radcliffe Hospital, Oxford, OX, DU, UK

^119^John P. Hussman Institute for Human Genomics, Leonard M. Miller School of Medicine, University of Miami, Miami, FL, USA

^120^The Dr. John T. Macdonald Foundation Department of Human Genetics, Leonard M. Miller School of Medicine, University of Miami, Miami, FL, USA

^121^F. Widjaja Foundation Inflammatory Bowel and Immunobiology Research Institute Cedars-Sinai Medical Center, Los Angeles, CA, USA

^122^Atherogenomics Laboratory University of Ottawa, Heart Institute, Ottawa, Canada

^123^Division of General Internal Medicine, Massachusetts General Hospital, Boston, MA, USA

^124^Department of Clinical Sciences University, Hospital Malmo Clinical Research Center, Lund University, Malmö, Sweden

^125^Estonian Genome Center, Institute of Genomics, University of Tartu, Tartu, Estonia

^126^University of Arizona Health Science, Tuscon, AZ, USA

^127^Department of Medicine, University of Maryland School of Medicine, Baltimore, MD, USA

^128^Howard Hughes Medical Institute and Department of Molecular Biology, Massachusetts General Hospital, Boston, MA, USA

^129^International Centre for Diarrhoeal Disease Research, Bangladesh

^130^Perelman School of Medicine, University of Pennsylvania, Philadelphia, PA, USA

^131^Johns Hopkins Bloomberg School of Public Health, Baltimore, MD, USA

^132^Lund University, Dept. Clinical Sciences, Skåne University Hospital, Malmö, Sweden

^133^Department of Statistical Genetics, Osaka University Graduate School of Medicine, Suita, Japan

^134^Laboratory of Statistical Immunology, Immunology Frontier Research Center (WPI-IFReC), Osaka University, Suita, Japan

^135^Integrated Frontier Research for Medical Science Division, Institute for Open and Transdisciplinary Research Initiatives, Osaka University, Suita, Japan

^136^Instituto Nacional de Medicina Genómica, (INMEGEN) Mexico City, Mexico

^137^Medical Research Institute, Ninewells Hospital and Medical School University of Dundee, Dundee, UK

^138^Wake Forest School of Medicine, Winston-Salem, NC, USA

^139^Department of Molecular Medicine and Biopharmaceutical Sciences, Graduate School of Convergence Science and Technology, Seoul National University, Seoul, Republic of Korea

^140^Department of Psychiatry Keck School of Medicine at the University of Southern California, Los Angeles, CA, USA

^141^Department of Psychiatry and Behavioral Sciences, Johns Hopkins University School of Medicine, Baltimore, MD, USA

^142^Children’s Hospital of Philadelphia, Philadelphia, PA, USA

^143^Division of Genetics and Epidemiology, Institute of Cancer Research, London, SM, NG

^144^University of Washington, Seattle, WA, USA

^145^Fred Hutchinson Cancer Research Center, Seattle, WA, USA

^146^Medical Research Center, Oulu University Hospital, Oulu Finland and Research Unit of Clinical Neuroscience Neurology University of Oulu, Oulu, Finland

^147^Center for Public Health Genomics, University of Virginia, Charlottesville, VA, USA

^148^Department of Public Health Sciences, University of Virginia, Charlottesville, VA, USA

^149^Research Center Montreal Heart Institute, Montreal, Quebec, Canada

^150^Department of Medicine, Faculty of Medicine Université de Montréal, Québec, Canada

^151^Department of Public Health Faculty of Medicine, University of Helsinki, Helsinki, Finland

^152^Broad Institute of MIT and Harvard, Cambridge, MA, USA

^153^Department of Biomedical Informatics Vanderbilt, University Medical Center, Nashville, TN, USA

^154^Department of Medicine, Vanderbilt University Medical Center, Nashville, TN, USA

^155^The Institute for Translational Genomics and Population Sciences, Department of Pediatrics, The Lundquist Institute for Biomedical Innovation at Harbor-UCLA Medical Center, Torrance, CA, USA

^156^Department of Biostatistics and Epidemiology, Perelman School of Medicine, University of Pennsylvania, Philadelphia, PA, USA

^157^Department of Medicine, Perelman School of Medicine at the University of Pennsylvania, Philadelphia, PA, USA

^158^Center for Non-Communicable Diseases, Karachi, Pakistan

^159^National Institute for Health and Welfare, Helsinki, Finland

^160^Deutsches Herzzentrum, München, Germany

^161^Technische Universität München, Germany

^162^Institute of Genetic Epidemiology, Department of Genetics and Pharmacology, Medical University of Innsbruck, 6020 Innsbruck, Austria

^163^Duke Molecular Physiology Institute, Durham, NC

^164^Division of Cardiovascular Medicine, Nashville VA Medical Center, Vanderbilt University School of Medicine, Nashville, TN, USA

^165^Division of Endocrinology, National University Hospital, Singapore

^166^NUS Saw Swee Hock School of Public Health, Singapore

^167^Channing Division of Network Medicine, Brigham and Women’s Hospital, Boston, MA, USA

^168^Harvard Medical School, Boston, MA, USA

^169^Department of Psychiatry, Icahn School of Medicine at Mount Sinai, New York, NY, USA

^170^Department of Genetics and Genomic Sciences, Icahn School of Medicine at Mount Sinai, New York, NY, USA

^171^Institute for Genomics and Multiscale Biology, Icahn School of Medicine at Mount Sinai, New York, NY, USA

^172^The Wallenberg Laboratory/Department of Molecular and Clinical Medicine, Institute of Medicine, Gothenburg University and the Department of Cardiology, Sahlgrenska University Hospital, Gothenburg, Sweden

^173^Department of Cardiology, Wallenberg Center for Molecular Medicine and Lund University Diabetes Center, Clinical Sciences, Lund University and Skåne University Hospital, Lund, Sweden

^174^Institute of Clinical Medicine Neurology, University of Eastern Finad, Kuopio, Finland

^175^Sorbonne Université, INSERM, Centre de Recherche Saint-Antoine, CRSA, AP-HP, Saint Antoine Hospital, Gastroenterology department, F-75012 Paris, France

^176^INRA, UMR1319 Micalis & AgroParisTech, Jouy en Josas, France

^177^Paris Center for Microbiome Medicine, (PaCeMM) FHU, Paris, France

^178^Department of Twin Research and Genetic Epidemiology King’s College London, London, UK

^179^The McDonnell Genome Institute at Washington University, Seattle, WA, USA

^180^Departments of Genetics and Psychiatry, University of North Carolina, Chapel Hill, NC, USA

^181^Saw Swee Hock School of Public Health National University of Singapore, National University Health System, Singapore

^182^Department of Medicine, Yong Loo Lin School of Medicine National University of Singapore, Singapore

^183^Duke-NUS Graduate Medical School, Singapore

^184^Life Sciences Institute, National University of Singapore, Singapore

^185^Department of Statistics and Applied Probability, National University of Singapore, Singapore

^186^Center for Behavioral Genomics, Department of Psychiatry, University of California, San Diego, CA, USA

^187^Institute of Genomic Medicine, University of California San Diego, San Diego, CA, USA

^188^Endocrinology, Abdominal Center, Helsinki University Hospital, Helsinki, Finland

^189^Institute of Genetics, Folkhalsan Research Center, Helsinki, Finland

^190^Juliet Keidan Institute of Pediatric Gastroenterology Shaare Zedek Medical Center, The Hebrew University of Jerusalem, Jerusalem, Israel

^191^Instituto de Investigaciones Biomédicas, UNAM, Mexico City, Mexico

^192^Instituto Nacional de Ciencias Médicas y Nutrición Salvador Zubirán, Mexico City, Mexico

^193^Department of Public Health Faculty of Medicine University of Helsinki, Helsinki, Finland

^194^Department of Psychiatry and Human Behavior, University of California Irvine, Irvine, CA, USA

^195^National Heart & Lung Institute & MRC London Institute of Medical Sciences, Imperial College, London, UK

^196^Cardiovascular Research Centre Royal Brompton & Harefield Hospitals, London, UK

^197^Radcliffe Department of Medicine, University of Oxford, Oxford, UK

^198^Department of Gastroenterology and Hepatology, University of Groningen and University Medical Center Groningen, Groningen, Netherlands

^199^Folkhälsan Institute of Genetics, Folkhälsan Research Center, Helsinki, Finland

^200^National Heart & Lung Institute and MRC London Institute of Medical Sciences, Imperial College London, London, UK

^201^Cardiovascular Research Centre, Royal Brompton & Harefield Hospitals NHS Trust, London, UK

^202^Department of Physiology and Biophysics, University of Mississippi Medical Center, Jackson, MS, USA

^203^Program in Infectious Disease and Microbiome, Broad Institute of MIT and Harvard, Cambridge, MA, USA

^204^Center for Computational and Integrative Biology, Massachusetts General Hospital, Boston, MA, USA

^205^Center for Genetic Epidemiology, Department of Population and Public Health Sciences, University of Southern California, Los Angeles, CA, USA

## Authors received funding as follows

Matthew J. Bown: British Heart Foundation awards CS/14/2/30841 and RG/18/10/33842

Josée Dupuis: National Heart Lung and Blood Institute’s Framingham Heart Study Contract (HHSNI); National Institute for Diabetes and Digestive and Kidney Diseases (NIDDK) R DK

Martti Färkkilä: State funding for university level health research

Laura D. Gauthier: Intel, Illumina

Stephen J. Glatt: U.S. NIMH Grant R MH

Leif Groop: The Academy of Finland and University of Helsinki: Center of Excellence for Complex Disease Genetics (grant number 312063 and 336822), Sigrid Jusélius Foundation; IMI 2 (grant No 115974 and 15881)

Mikko Hiltunen: Academy of Finland (grant 338182) Sigrid Jusélius Foundation the Strategic Neuroscience Funding of the University of Eastern Finland

Chaim Jalas: Bonei Olam

Ronald Ma: Research Grants Council Theme-based Research Scheme (T12-402/13N), RGC Research Impact Fund (CU R4012-18) and a Croucher Foundation Senior Medical Research Fellowship.

Jaakko Kaprio: Academy of Finland (grants 312073 and 336823)

Jacob McCauley: National Institute of Diabetes and Digestive and Kidney Disease Grant R01DK104844

Yukinori Okada: JSPS KAKENHI (19H01021, 20K21834), AMED (JP21km0405211, JP21ek0109413, JP21gm4010006, JP21km0405217, JP21ek0410075), JST Moonshot R&D (JPMJMS2021)

Michael J. Owen: Medical Research Council UK: Centre Grant No. MR/L010305/1, Program Grant No. G0800509 Aarno Palotie: the Academy of Finland Center of Excellence for Complex Disease Genetics (grant numbers 312074 and 336824) and Sigrid Jusélius Foundation

John D. Rioux: National Institute of Diabetes and Digestive and Kidney Diseases (NIDDK; DK062432), from the Canadian Institutes of Health (CIHR GPG 102170), from Genome Canada/Génome Québec (GPH-129341), and a Canada Research Chair (#230625)

Samuli Ripatti: the Academy of Finland Center of Excellence for Complex Disease Genetics (grant number) Sigrid

## Jusélius Foundation

Jerome I. Rotter: Trans-Omics in Precision Medicine (TOPMed) program was supported by the National Heart, Lung and Blood Institute (NHLBI). WGS for “NHLBI TOPMed: Multi-Ethnic Study of Atherosclerosis (MESA)” (phs001416.v1.p1) was performed at the Broad Institute of MIT and Harvard (3U54HG003067-13S1). Core support including centralized genomic read mapping and genotype calling, along with variant quality metrics and filtering were provided by the TOPMed Informatics Research Center (3R01HL-117626-02S1; contract HHSN268201800002I). Core support including phenotype harmonization, data management, sample-identity QC, and general program coordination were provided by the TOPMed Data Coordinating Center (R01HL-120393; U01HL-120393; contract HHSN268201800001I). We gratefully acknowledge the studies and participants who provided biological samples and data for MESA and TOPMed. JSK was supported by the Pulmonary Fibrosis Foundation Scholars Award and grant K23-HL-150301 from the NHLBI. MRA was supported by grant K23-HL- 150280, AJP was supported by grant K23-HL-140199, and AM was supported by R01-HL131565 from the NHLBI. EJB was supported by grant K23-AR-075112 from the National Institute of Arthritis and Musculoskeletal and Skin Diseases.The MESA project is conducted and supported by the National Heart, Lung, and Blood Institute (NHLBI) in collaboration with MESA investigators. Support for MESA is provided by contracts 75N92020D00001, HHSN268201500003I, N01-HC-95159, 75N92020D00005, N01-HC-95160, 75N92020D00002, N01-HC-95161, 75N92020D00003, N01-HC-95162, 75N92020D00006, N01-HC-95163, 75N92020D00004, N01-HC-95164, 75N92020D00007, N01-HC-95165, N01-HC-95166, N01-HC-95167, N01-HC-95168, N01-HC-95169, UL1-TR-000040, UL1-TR-001079, and UL1-TR-001420. Also supported in part by the National Center for Advancing Translational Sciences, CTSI grant UL1TR001881, and the National Institute of Diabetes and Digestive and Kidney Disease Diabetes Research Center (DRC) grant DK063491 to the Southern California Diabetes Endocrinology Research Center

Edwin K. Silverman: NIH Grants U01 HL089856 and U01 HL089897

J. Gustav Smith: The Swedish Heart-Lung Foundation (2019-0526), the Swedish Research Council (2017-02554), the European Research Council (ERC-STG-2015-679242), Skåne University Hospital, governmental funding of clinical research within the Swedish National Health Service, a generous donation from the Knut and Alice Wallenberg foundation to the Wallenberg Center for Molecular Medicine in Lund, and funding from the Swedish Research Council (Linnaeus grant Dnr 349-2006-237, Strategic Research Area Exodiab Dnr 2009-1039) and Swedish Foundation for Strategic Research (Dnr IRC15-0067) to the Lund University Diabetes Center

Kent D. Taylor: Trans-Omics in Precision Medicine (TOPMed) program was supported by the National Heart, Lung and Blood Institute (NHLBI). WGS for “NHLBI TOPMed: Multi-Ethnic Study of Atherosclerosis (MESA)” (phs001416.v1.p1) was performed at the Broad Institute of MIT and Harvard (3U54HG003067-13S1). Core support including centralized genomic read mapping and genotype calling, along with variant quality metrics and filtering were provided by the TOPMed Informatics Research Center (3R01HL-117626-02S1; contract HHSN268201800002I). Core support including phenotype harmonization, data management, sample-identity QC, and general program coordination were provided by the TOPMed Data Coordinating Center (R01HL-120393; U01HL-120393; contract HHSN268201800001I). We gratefully acknowledge the studies and participants who provided biological samples and data for MESA and TOPMed. JSK was supported by the Pulmonary Fibrosis Foundation Scholars Award and grant K23-HL-150301 from the NHLBI. MRA was supported by grant K23-HL- 150280, AJP was supported by grant K23-HL-140199, and AM was supported by R01-HL131565 from the NHLBI. EJB was supported by grant K23-AR-075112 from the National Institute of Arthritis and Musculoskeletal and Skin Diseases.The MESA project is conducted and supported by the National Heart, Lung, and Blood Institute (NHLBI) in collaboration with MESA investigators. Support for MESA is provided by contracts 75N92020D00001, HHSN268201500003I, N01-HC-95159, 75N92020D00005, N01-HC-95160, 75N92020D00002, N01-HC-95161, 75N92020D00003, N01-HC-95162, 75N92020D00006, N01-HC-95163, 75N92020D00004, N01-HC-95164, 75N92020D00007, N01-HC-95165, N01-HC-95166, N01-HC-95167, N01-HC-95168, N01-HC-95169, UL1-TR-000040, UL1-TR-001079, and UL1-TR-001420. Also supported in part by the National Center for Advancing Translational Sciences, CTSI grant UL1TR001881, and the National Institute of Diabetes and Digestive and Kidney Disease Diabetes Research Center (DRC) grant DK063491 to the Southern California Diabetes Endocrinology Research Center

Tiinamaija Tuomi: The Academy of Finland and University of Helsinki: Center of Excellence for Complex Disease Genetics (grant number 312072 and 336826), Folkhalsan Research Foundation, Helsinki University Hospital, Ollqvist Foundation, Liv och Halsa foundation; NovoNordisk Foundation

Teresa Tusie-Luna: CONACyT Project 312688

James S. Ware: Sir Jules Thorn Charitable Trust [21JTA], Wellcome Trust [107469/Z/15/Z], Medical Research Council (UK), NIHR Imperial College Biomedical Research Centre

Rinse K. Weersma: The Lifelines Biobank initiative has been made possible by subsidy from the Dutch Ministry of Health Welfare and Sport the Dutch Ministry of Economic Affairs the University Medical Centre Groningen (UMCG the Netherlands) the University of Groningen and the Northern Provinces of the Netherlands

No conflicts of interest to declare.

## Notes

### Competing Interest Statement

The authors have declared no competing interest.

### Summary of Updates

Constraint model (& scores) updated with extended validation analyses

