## Supplemental Information for "A genome-wide mutational constraint map quantified from variation in 76,156 human genomes"

**Supplemental Information** 1

**Data processing and variant calling** 3

Supplementary Figure 1 | Growth of data size with number of samples. 4

**Sample QC** 5

Hard filtering 5

Sex inference 5

Supplementary Table 1 | Sex chromosome inference. 6

Supplementary Figure 2 | Sex inference. 6

Defining a high quality set of sites for QC 7

Relatedness inference 7

Ancestry assignment 7

Supplementary Figure 3 | UMAP embedding of PCA results (PCs 1-6 and 8-16) of all individuals in the gnomAD release. 8

Filtering based on QC metrics 8

Supplementary Figure 4 | The distribution of the number of SNPs for the African ancestry samples using the clustering-based and regression-based approach. 9

Supplementary Table 2 | Final number of individuals for each population. 10

**Variant QC and annotation** 10

Supplementary Figure 5 | Precision-recall curves for the previous site-level (VQSR), allele-specific (AS_VQSR) and allele-specific with transmitted singletons (AS_VQSR_TS) approaches. 11

Functional annotation 12

**Constraint modeling and assessment** 12

Estimation of trinucleotide context-specific mutation rates 12

Supplementary Figure 6 | Trinucleotide context-specific mutation rate estimates. 13

Comparison of constraint Z score and other predictive scores 13

Supplementary Figure 7 | Performance of different metrics in identifying putative functional variants in stringent non-coding regions. 14

Analysis of constraint Z scores for chromosome X 15

Supplementary Figure 8 | Distribution of constraint Z scores on chromosome X. 15

Power of constraint detection 16

Supplementary Figure 9 | Power analysis. 18

**Code and data availability** **18**

Release files 18

Code availability 19

The gnomAD browser 19

Support for multiple reference genomes 19

Supplementary Figure 10 | The same gene viewed in gnomAD v2 and v3. 19

Supplementary Figure 11 | Liftover section on a gnomAD v2 and v3 variant page. 19

HGDP and 1000 Genomes population frequencies 20

Supplementary Figure 12 | Detailed population frequencies. Here, we show the frequency table for the 1000 Genomes project. 20

Read data in non-coding regions 20

Supplementary Datasets 21

**References** **22**

The version of the Genome Aggregation Database (gnomAD) v3 presented in this manuscript is a catalog containing 759,302,267 short nuclear variants (644,267,978 passing stringent variant quality control [QC]) based on whole-genome sequencing of 76,156 samples (passing QC from an initial collection of 153,030 samples) mapped to the GRCh38 build of the human reference genome. In this release, we have included more than 3,000 new samples specifically chosen to increase the ancestral diversity of the resource, and for the first time, we provide individual genotypes in addition to variant calls for a subset of gnomAD, which includes new data from >60 distinct populations from Africa, Europe, the Middle East, South and Central Asia, East Asia, Oceania, and the Americas. Many of the processing, quality control, and analysis procedures closely resemble those from the 15,748 genomes from the gnomAD v2 manuscript (Karczewski et al. 2020). In this supplement, we highlight the differences where applicable.

**Data processing and variant calling**

Whole genome sequences were mapped using bwa mem 0.7.15.r1140 against the GRCh38 version hs38DH, which includes decoy contigs and HLA genes (FASTA located at https://console.cloud.google.com/storage/browser/gcp-public-data--broad-references/hg38/v0/). Reads were then processed using the GATK best practices using GATK4 for BQSR and GATK3.5 for HaplotypeCaller to produce gVCFs.

Previous approaches (e.g. gnomAD v2) have typically involved the joint calling of the full cohort using GATK to produce a VCF file with a genotype for each sample at every position where at least one sample contains a non-reference allele. However, this approach would not scale to 150,000 samples, due to time and memory limitations, as well as storage: the output would require about 900TB and be prohibitively expensive to store and compute over.

Instead, we implemented and used a novel combiner within Hail (described in detail in (Karczewski et al. 2021)), which combines gVCFs into a sparse MatrixTable (MT). gVCFs are single-sample files, which contain one row for each genomic position where a non-reference allele is found in the sample and, unlike VCFs, a row for each reference block start. Reference blocks are contiguous bases where the sample is homozygous reference within certain confidence boundaries. In gnomAD v3, we used the following three confidence bins: No coverage / evidence; Genotype quality < Q20; and Genotype quality >= Q20. For each of these bins, the reference block stores the minimum and median coverage, and the minimum genotype quality for the bases residing in the block.

Using this new sparse data format, the full gnomAD v3 MT only requires 20TB of storage (Supplementary Figure 1). This new format scales linearly with the number of samples and is lossless with respect to the input sample gVCFs. Importantly, much more granular QC metrics, previously collapsed into the INFO field across samples, are preserved at each non-reference genotype in the data, such as strand balance, read position metrics (ReadPosRankSum), etc. Thus, new data can be appended to existing data without re-processing of the previously processed samples. We demonstrated the power of this data format by adding 4,598 genomes to our original gnomAD v3 release of 71,702 genomes. Finally, while not currently implemented, it is possible to re-export a gVCF from this format, removing the need for storing the gVCFs.

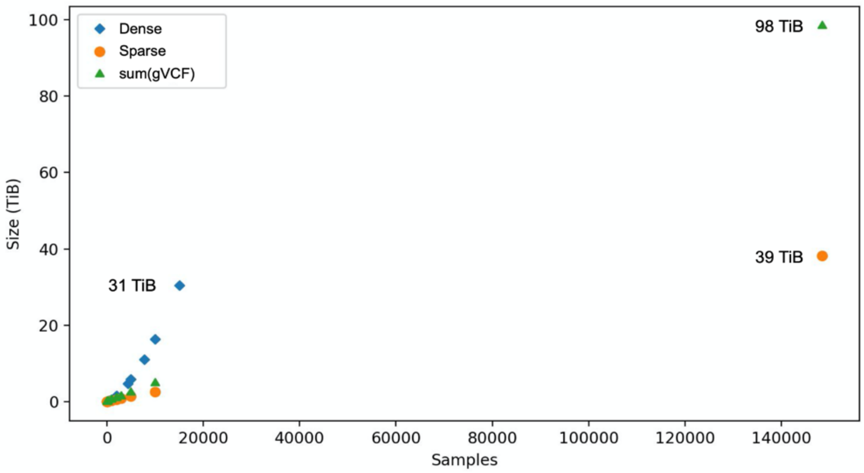


**Supplementary Figure 1 | Growth of data size with number of samples.**
The dense representation (VCF) grows super-linearly, while the aggregate gVCF size and the SparseMT representation grow linearly. The final gnomAD v3 sparse dataset was smaller still (20TiB) due to increased compression from broader reference block confidence bins.

### Sample QC

The sample QC process was similar to that of gnomAD v2 (Karczewski et al. 2020). Briefly, hard filters were applied to remove samples of poor quality as well as samples that did not have permissions for public release of aggregate data. Next, we inferred sex for each sample and removed samples with sex chromosome aneuploidies or ambiguous sex assignment: here, we modified the original pipeline by using normalized coverage on both X and Y in order to infer sample sex. We defined and used a set of high quality sites to infer relatedness between samples, allowing us to filter to a set of unrelated individuals, and assign ancestry to each sample. Finally, we filtered samples that were determined to be outliers based on sample QC metrics, using a novel regression-based method. All quality control and processing steps were performed using Hail 0.2.62 (Hail Team. Hail 0.2.62-84fa81b9ea3d. https://github.com/hail-is/hail/commit/84fa81b9ea3d.).

#### Hard filtering

We computed sample QC metrics using the Hail 'sample_qc' module on all autosomal bi-allelic single nucleotide variants (SNVs). We removed samples that were clear outliers for the number of SNVs (< 2.4 million or > 3.75 million), number of singletons (> 100,000), ratio of heterozygous to homozygous variants > 3.3, and a mean coverage on chromosome 20 of < 15X. Additionally, for 87,756 of the 92,306 releasable samples where BAM-level metrics were available, we removed samples that were outliers for percent contamination (> 5%), percent chimeras: (> 5%), and median insert size (< 250bp).

#### Sex inference

To infer sex, we computed the mean coverage on non-pseudoautosomal (non-PAR) regions of chromosome X and Y and normalized these values using the mean coverage on chromosome 20. In addition, we ran the Hail 'impute_sex' function on the non-PAR regions of chromosome X to compute the inbreeding coefficient F-stat. Based on these three metrics, we assigned a number of X chromosomes based on normalized X coverage 0-1.2913 (1 X), 1.4477-2.3961 (2 X), or 2.4909+ (3 X), and a number of Y chromosomes based on normalized Y coverage 0-0.1 (no Y), 0.1-1.1645 (1 Y), 1.2381+ (2 Y). The final assignments are shown in Supplementary Table 1 and Supplementary Figure 2.

**Supplementary Table 1 | Sex chromosome inference.**
The coverages for normalized X and Y coverages are shown for each sex chromosome inference assignment, alongside total number of releasable samples that pass QC to this point.

| | | **X chromosome coverage** | | **Y chromosome coverage** | |
|---|---|---|---|---|---|
| **Inferred sex chromosomes** | **Total** | **Lower cutoff** | **Upper cutoff** | **Lower cutoff** | **Upper cutoff** |
| XX | 46,361 | 1.4477 | 2.3961 | 0 | 0.1 |
| XY | 45,129 | 0 | 1.2913 | 0.1 | 1.1645 |
| XO | 425 | 0 | 1.2913 | 0 | 0.1 |
| XXY | 107 | 1.4477 | 2.3961 | 1.2381 | 1.1645 |
| XXX | 34 | 2.4909 | - | 0 | 0.1 |
| XYY | 31 | 0 | 1.2913 | 1.2381 | - |
| XXXY | 4 | 2.4909 | - | 0.1 | 1.1645 |
| XXYY | 3 | 1.4477 | 2.3961 | 1.2381 | - |
| ambiguous | 212 | All others | | All others | |

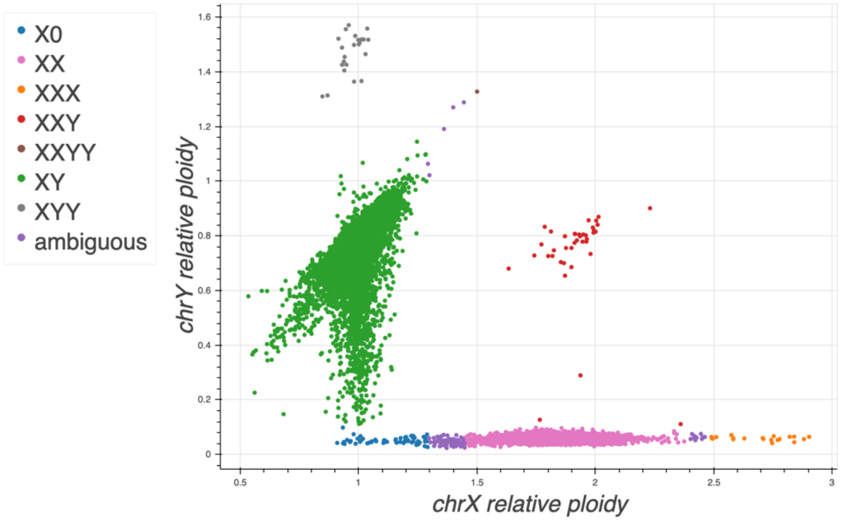


**Supplementary Figure 2 | Sex inference.**
A scatter plot of ploidy on chromosomes X and Y is shown for each individual in the dataset. Points are colored by inferred sex haplotype.

#### Defining a high quality set of sites for QC

In order to perform relatedness and ancestry inference, we first selected a set of high quality QC sites as follows:

1. We took all sites that were used for gnomAD v2.1 and lifted them over to GRCh38
2. We added ~5k sites widely used for quality control of GWAS data (Purcell et al. 2014) and lifted these sites over to GRCh38
3. From these two sets of sites, we then selected all bi-allelic SNVs with an Inbreeding coefficient > -0.25 (no excess of heterozygotes)

In total, we ended up with 76,419 high quality variants for relatedness and ancestry inference.

#### Relatedness inference

We used PC-Relate (implemented in Hail 'pc_relate') (Conomos et al. 2016) to compute relatedness, followed by Hail's 'maximal_independent_set' in order to select as many samples as possible, while asserting that the final dataset includes no pairs of first and second degree relatives. When multiple samples could be selected, we kept the sample with the highest coverage as a tie-breaker.

#### Ancestry assignment

We used principal component analysis (PCA; using the 'hwe_normalized_pca' function in Hail) on the set of high quality variants in our unrelated samples, and selected the first 16 PCs to assign ancestry. We then trained a random forest classifier using 22,054 samples with known ancestry and 14,828 samples for which we had a population label from gnomAD v2 as training samples and using the PCs as features. We assigned ancestry to all samples for which the probability of that ancestry was > 75% according to the random forest model. All other samples were unassigned (labeled oth). A UMAP embedding of the first

PCs is shown in Supplementary Figure 3 (Diaz-Papkovich, Anderson-Trocme, and Gravel 2018; McInnes et al. 2018).

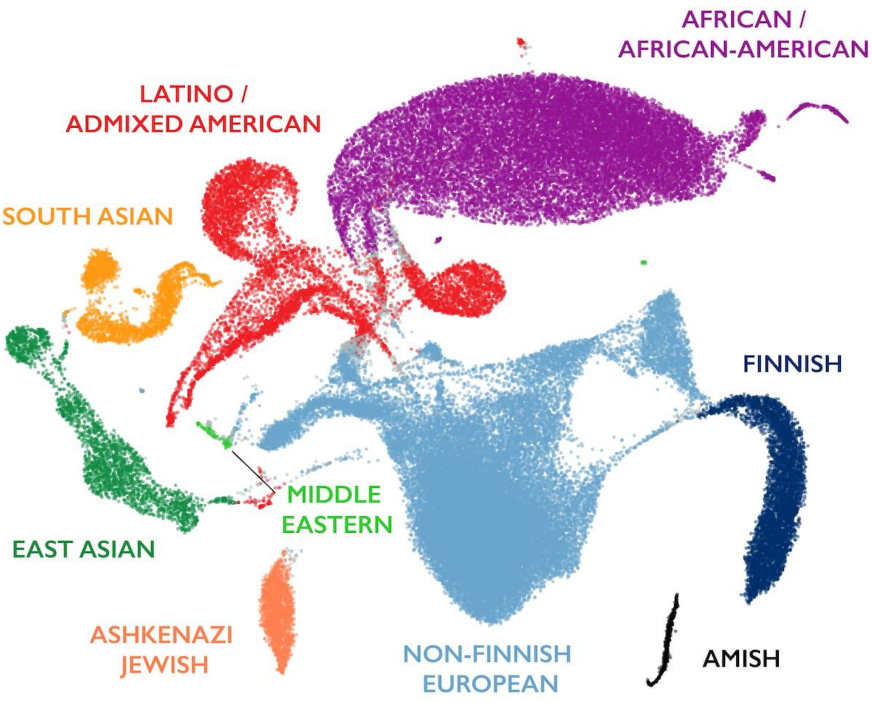


**Supplementary Figure 3 | UMAP embedding of PCA results (PCs 1-6 and 8-16) of all individuals in the gnomAD release.**
This embedding is an illustration of the structure present in the dataset: note that long-range distances in this projection do not reflect genetic distance between populations.

#### Filtering based on QC metrics

In gnomAD v2, we grouped samples based on their ancestry assignments and filtered outliers within each ancestry based on various quality metrics such as number of SNPs. Here, we built a single model for each metric across all individuals, regressing out the PCs computed during the ancestry assignment, and filtered samples based on the residuals for each of the QC metrics. This strategy allowed us to consider the samples' ancestry as a continuous spectrum and was particularly beneficial for admixed samples and samples that did not get an ancestry assignment (assigned as oth; previously, we lumped all these samples together even though they were drawn from multiple populations). An illustration of the improvement from this new method is shown in Supplementary Figure 4.

**Clustering-based approach** **Regression-based approach**

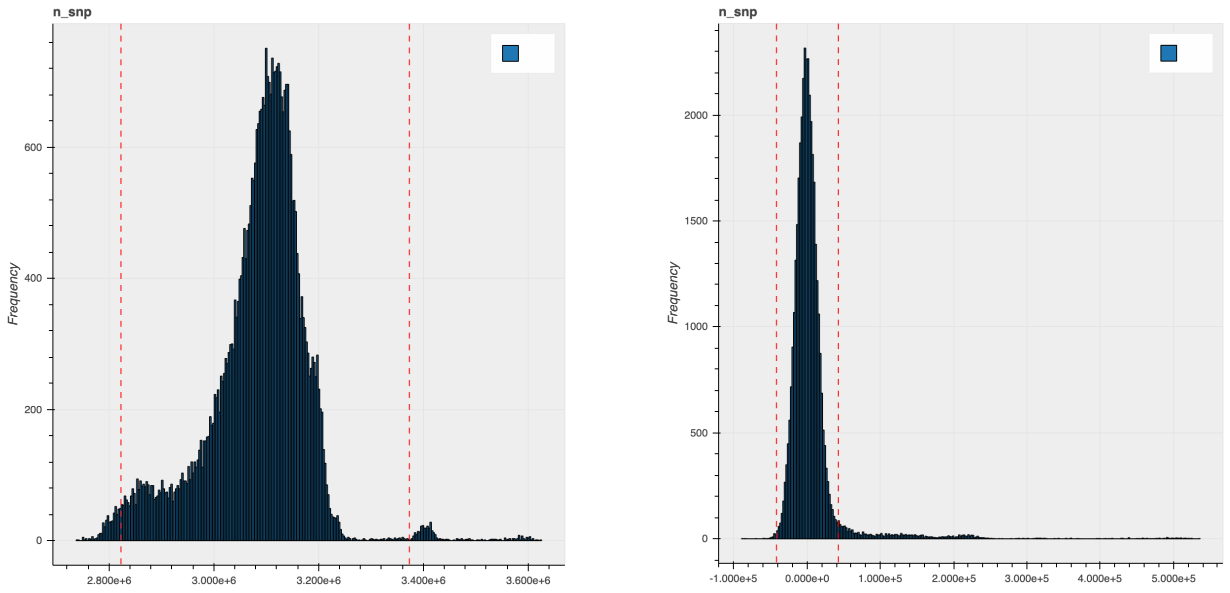


**Supplementary Figure 4 | The distribution of the number of SNPs for the African ancestry samples using the clustering-based and regression-based approach.**

We used the sample QC metrics computed using the Hail 'sample_qc' module on all autosomal bi-allelic SNVs and filtered samples that were 4 median absolute deviations (MADs) from the median for the following metrics: n_snp, r_ti_tv, r_insertion_deletion, n_insertion, n_deletion, r_het_hom_var, n_het, n_hom_var, n_transition and n_transversion. In addition, we filtered samples that fell outside 8 MADs above the median n_singleton metric and over 4 MADs above the median r_het_hom_var metric. The final set of samples is shown in Supplementary Table 2.

After some downstream analysis, we noted that this regression approach removes samples from populations with extreme diversity, such as individuals from the San, Papuan, and Pygmy populations in HGDP. We have adjusted this filter for future releases and provide the data for all these individuals in a joint-called subset of gnomAD (see Code and Data Availability section below, as well as accompanying manuscript) under release v3.1.2. However, we note that this current dataset excludes these individuals, as newly computing the frequency metrics was prohibitively expensive.

**Supplementary Table 2 | Final number of individuals for each population.**

| Population code | Description | Number of Genomes |
|---|---|---|
| afr | African/African American | 20,744 |
| ami | Amish | 456 |
| amr | Latino/Admixed American | 7,647 |
| asj | Ashkenazi Jewish | 1,736 |
| eas | East Asian | 2,604 |
| fin | Finnish | 5,316 |
| nfe | Non-Finnish European | 34,029 |
| mid | Middle Eastern | 158 |
| sas | South Asian | 2,419 |
| oth | Other (population not assigned) | 1,047 |
| Total | | 76,156 |

### Variant QC and annotation

Because the new sparse MatrixTable format contains all the information encoded in the gVCFs, we computed all variant QC metrics within Hail, which enabled the separate computation for each allele (rather than site-level as was typically done previously). The code to compute these metrics is available at https://github.com/broadinstitute/gnomad_methods/blob/master/gnomad/utils/sparse_mt.py. We then used the allele-specific version of GATK Variant Quality Score Recalibration (VQSR) to compute a confidence score for each allele in the dataset. We used the following allele-specific features: FS, SOR, ReadPosRankSum, MQRankSum, and QD for SNPs and indels, as well as MQ for SNPs.

In addition to the GATK bundle training resources (HapMap, Omni, 1000 Genomes, and Mills indels), we also used a set of ~19M transmitted singletons (alleles observed exactly twice in the dataset, only in a parent/child duo) from 6,743 trios present in our raw data.

We assessed the results of the filtering by plotting, as a function of quality tranche, the number of potential *de novo* mutations in the 6,743 trios, the Ti/Tv ratio, proportion singletons, proportion bi-allelic variants, and variants in ClinVar, as well as precision and recall in two truth samples present in our data: NA12878 and a pseudo-diploid sample (A mixture of DNA [est. 50.7% / 49.3%] from two haploid CHM cell lines).

In gnomAD v3 (prior to the addition of 4,598 new samples), we assessed the performance of the classic site-level VQSR, to a new algorithm, the allele-specific VQSR (AS_VQSR), as well as one with transmitted singletons included (AS_VQSR_TS). Supplementary Figure 5 illustrates the superior performance of the allele-specific approach by precision-recall curves of gold standard SNVs and indels from one sample, NA12878.

**NA12878 SNVs** **NA12878 Indels**

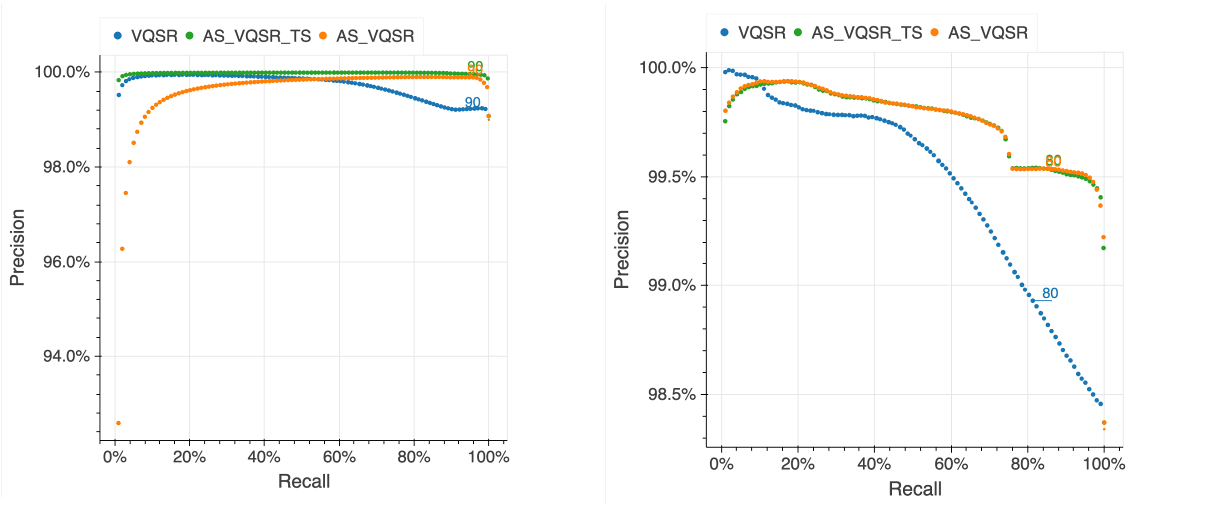


**Supplementary Figure 5 | Precision-recall curves for the previous site-level (VQSR), allele-specific (AS_VQSR) and allele-specific with transmitted singletons (AS_VQSR_TS) approaches.**
Both allele-specific approaches outperform VQSR, while AS_VQSR_TS shows a slight improvement for SNVs.

In addition to VQSR, we also applied the following hard filters:

- AC0: No sample had a high quality genotype at this variant site (GQ>=20, DP>=10 and allele balance > 0.2 for heterozygotes)
- InbreedingCoeff: there was an excess of heterozygotes at the site compared to Hardy-Weinberg expectations using a threshold of -0.3 on the InbreedingCoefficient metric.

In total, 12.2% of SNVs and 32.5% of indels were filtered, resulting in 569,860,911 SNVs and 74,407,067 indels that passed all filters in our release.

##### Functional annotation

All variants are annotated using version 101 of the Variant Effect Predictor (VEP) based on the gene models from Gencode v35, with the LOFTEE plugin as described previously (Karczewski et al. 2020). For GRCh38, LOFTEE is similar to the previous implementation, without the extended splice predictions.

#### Constraint modeling and assessment

##### Estimation of trinucleotide context-specific mutation rates

To calculate the baseline mutation rate for each substitution in a trinucleotide context ($XY_1Z$ -> $XY_2Z$), we count the instances of each trinucleotide context in the autosomes of the human genome, excluding sites where 1) a low-quality variant is called in gnomAD v3 (INFO/FILTER does not equal to 'PASS'), 2) the mean coverage in the gnomAD genomes is <30X or >32X, or 3) the region is marked as 'blacklisted regions' (ENCODE https://www.encodeproject.org/files/ENCFF356LFX/) or 'gaps' (telomeres and centromeres). This resulted in 6,079,733,538 possible variants at 2,026,577,846 autosomal sites. Using this dataset, we compute the proportion of possible variants observed (hereafter referred to as 'proportion observed') for each substitution and context, using rare variants observed in gnomAD with an allele frequency (AF) ≤0.1%. Our estimates are well-correlated with the mutation rates reported by Carlson et al using an independent dataset (the Bipolar Research in Deep Genome and Epigenome Sequencing [BRIDGES] study; Supplementary Figure 6a) and are highly stable across different AF thresholds (0.01%-1%; Supplementary Figure 6b).

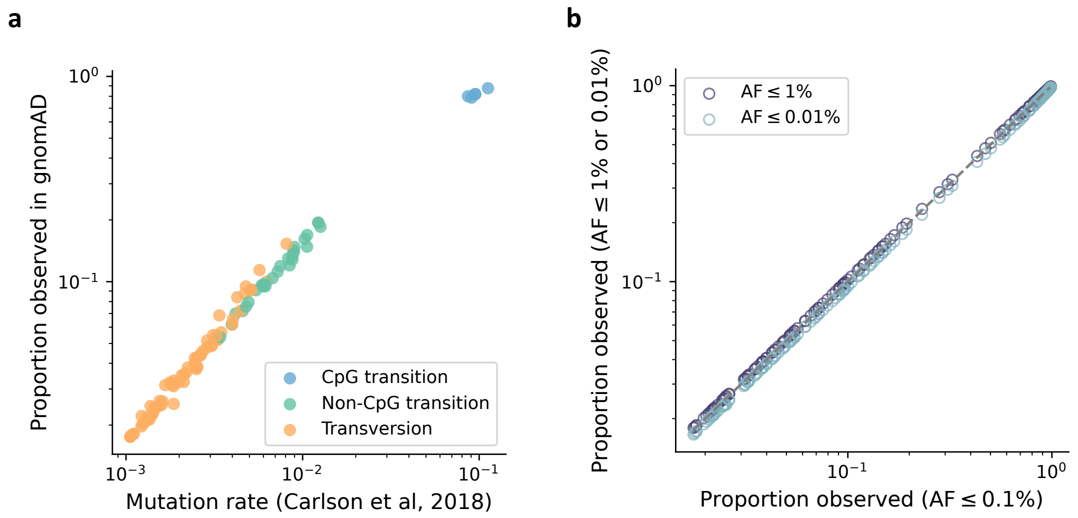


**Supplementary Figure 6 | Trinucleotide context-specific mutation rate estimates.**
The mutation rates estimated and used in this study are well-correlated with previous estimates from Carson et al using an independent dataset (a) and are highly stable across different AF thresholds (b).

#### Comparison of constraint Z score and other predictive scores

Benchmarking on various validation datasets, we show that our constraint Z score outperforms other constraint/conservation metrics – Orion, CDTS, gwRVIS, DR, phyloP, phastCons, and GERP – in identifying putative functional non-coding variants (Fig. 3 and Extended Fig. 4). We note that all evaluations were performed within the non-coding genome for explicitly comparing the metrics in prioritizing non-coding variants. We further eliminate potential bias from nearby genes by recapitulating the results within regions >10kb away from any protein-coding exons (Supplementary Fig. 7).

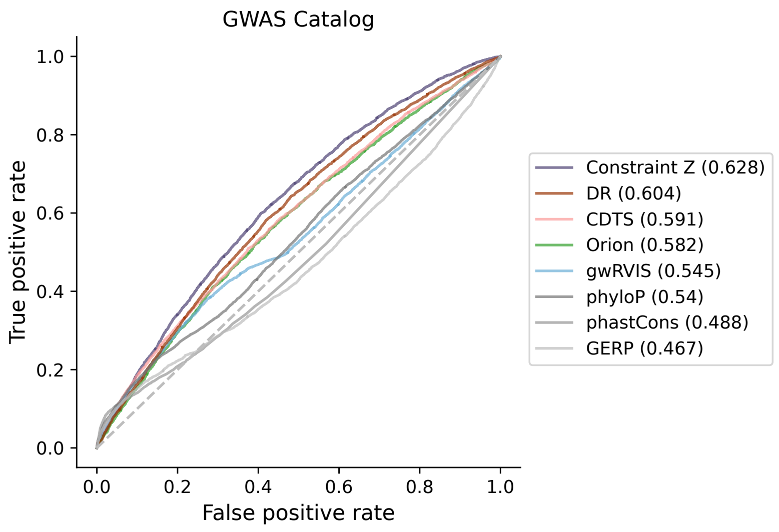


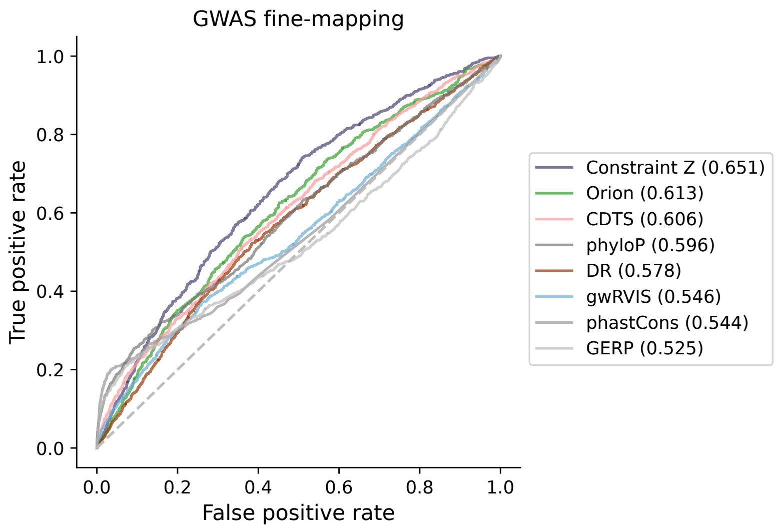


**Supplementary Figure 7 | Performance of different metrics in identifying putative functional variants in stringent non-coding regions.**

As described in Methods, the stringent non-coding variant sets include 4,379 GWAS Catalog variants, 967 GWAS fine-mapping variants, a high-confidence subset of 59 fine-mapped variants, and 7 ClinVar pathogenic variants. Analyses were not performed for the latter two due to small sample sizes.

#### Analysis of constraint Z scores for chromosome X

Due to the lack of DNM data on chromosome X, we built our mutational model using autosomal regions and extrapolated it to construct constraint Z scores for chromosome X. Consistent with the results in autosomes, protein-coding sequences overall show a significantly higher constraint Z score than non-coding regions (median=1.72 versus 0.36, Wilcoxon $P<10^{-200}$; Supplementary Fig. 8a), and constrained non-coding regions are significantly enriched for regulatory elements (e.g., ENCODE cCREs; Supplementary Fig. 8b).

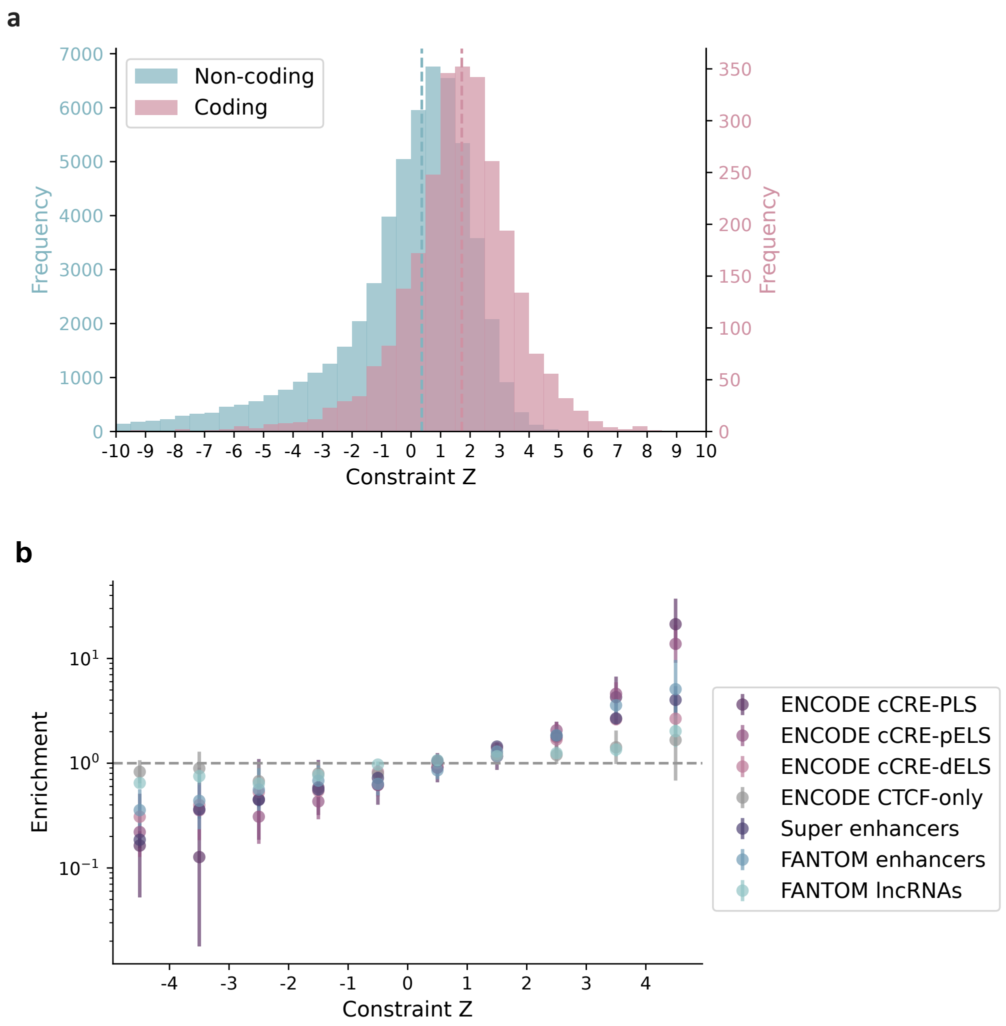


**Supplementary Figure 8 | Distribution of constraint Z scores on chromosome X.**
a, Windows overlapping coding regions (N=2,647 with ≥ 1bp coding sequence; red) overall exhibit a higher constraint Z (stronger negative selection) than windows that are exclusively non-coding (N=55,055; blue). Dashed

lines indicate the medians. b, Constrained non-coding regions are enriched for regulatory elements (see Methods for details in the collection of different annotations). Enrichment was evaluated by comparing the proportion of non-coding 1kb windows, binned by constraint Z, that overlap with a given functional annotation to the genome-wide average. Error bars indicate 95% confidence intervals of the odds ratios.

#### Power of constraint detection

We estimate the power of our metric in detecting constraint as the percentage of the non-coding genome to obtain a high constraint Z score (Z≥4) under a certain level of negative selection. We quantify the strength of negative selection in a region by the level of depletion of variation (1-observed/expected). For a given depletion of variation, the minimum number of expected variants to achieve a Z≥4 is determined, and the number of samples required to achieve the expected number of variants is estimated using a linear model of log(number of expected variants) ~ log(number of samples) from downsampling the gnomAD dataset. We benchmark the power by the depletion of variation observed in coding exons of similar size – at a 1kb scale, we are currently well-powered (≥90%) to detect extreme non-coding constraint as strong as the 90$^{th}$ percentile of coding exons (42% depletion), and we estimate a sample size of ~340K genomes to detect constraint as to the 50$^{th}$ percentile (20% depletion; Supplementary Fig. 9a). Much larger sample sizes will be needed for further increasing the resolution – at a 100bp scale, we would need ~5.3M samples to detect the 50$^{th}$ percentile (26% depletion; Supplementary Fig. 9b). Under the current sample size, we show that 1kb is the optimal window size by comparing it to other window sizes (100bp, 500bp, 2kb, and 3kb) in prioritizing non-coding variants (Supplementary Fig. 9c).

Meanwhile, we emphasize the importance of increasing ancestry diversity in the dataset to improve power. A more diverse population would allow more rare variants to be observed and thereby increase the power of detecting depletions of variation. We explicitly demonstrate this by reconstructing our constraint metric from the subset of European population (N=34,029) and comparing it to that from an equal-sized subset containing all diverse populations. The diversified subset is shown to achieve a higher predictive power in identifying functional non-coding variants (Supplementary Fig. 9d).

**a**

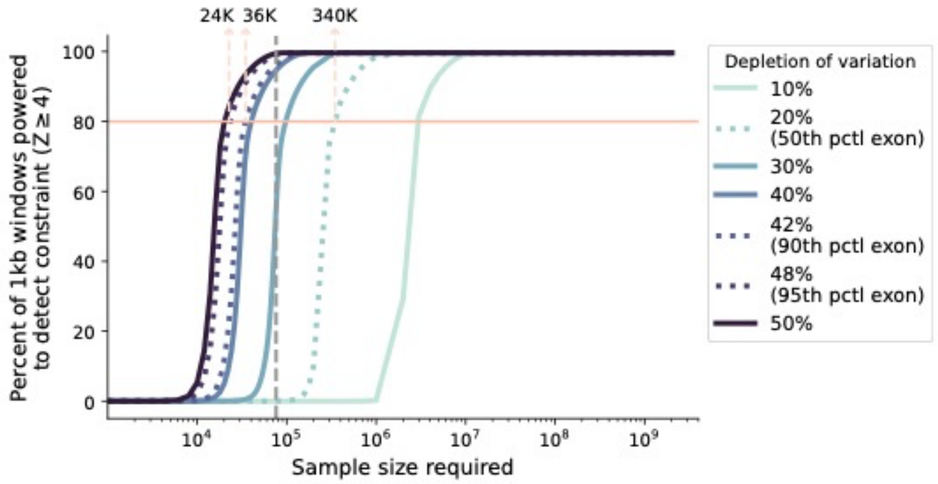


**b**

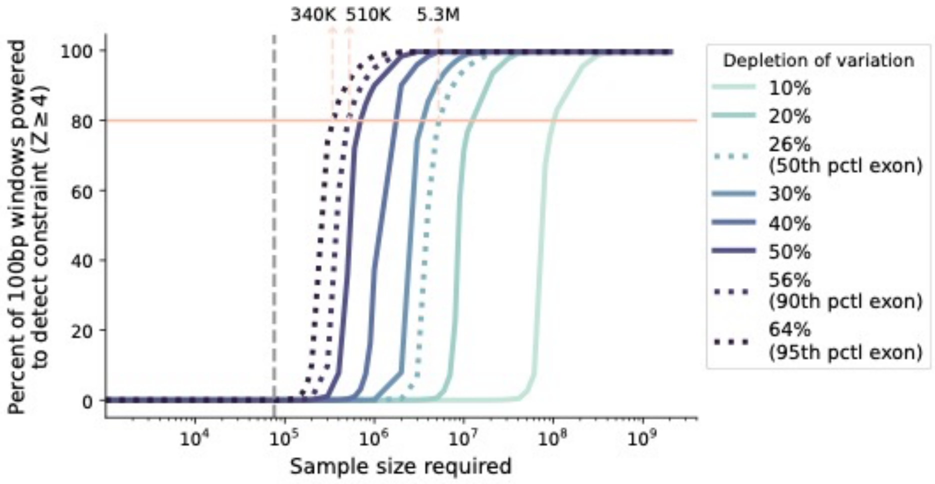

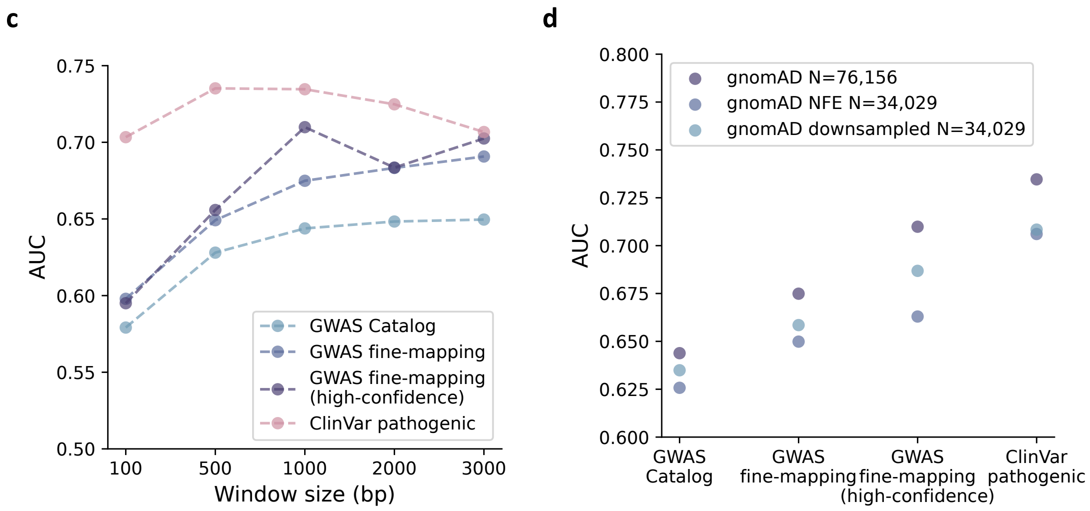


**Supplementary Figure 9 | Power analysis.**
a,b, The sample size required for well-powered non-coding constraint detection. The percentage of non-coding regions powered to detect constraint (Z≥4) at a 1kb (a) and 100bp (b) scale under varying levels of selection (depletion of variation) is shown as a function of log-scaled sample size. Lighter color indicates milder deletion of variation (weaker selection), which requires a larger sample size to detect constraint; the grey dashed vertical line indicates the current sample size of 76,156 genomes. Dotted curves (left to right) benchmark the 95$^{th}$, 90$^{th}$, and 50$^{th}$ percentile of depletion of variation observed in coding exons of similar size. The number of samples required to obtain 80% detection power is labeled at corresponding benchmarks. c, AUCs of constraint Z scores computed on different window sizes in identifying putative functional non-coding variants. 1kb (used in this study) presents the optimal window size with high performance while maintaining reasonable resolution. d, AUCs of constraint Z scores computed from different subsets of gnomAD in identifying putative functional non-coding variants. While with an equal sample size, the downsampled dataset with diverse ancestries presents better performance than the Non-Finnish European (NFE)-only dataset.

### Code and data availability

#### Release files

We release the aggregated allele frequency dataset at https://gnomad.broadinstitute.org, in a browser and bulk downloads for VCFs and Hail Tables, as well as all constraint statistics described in this manuscript. Additionally, we provide a subset of the dataset that includes individual level data for the HGDP (Bergström et al. 2020) and the 1000 Genomes projects (1000 Genomes Project Consortium et al. 2015): the generation and use of this dataset is described in a companion manuscript.

#### Code availability

All code to perform quality control of the resource is publicly available at https://github.com/broadinstitute/gnomad_qc, and many of the functions are documented in a Python package (gnomad) at https://broadinstitute.github.io/gnomad_methods/index.html. The code to compute the constraint statistics is available at https://github.com/atgu/gnomad_nc_constraint.

#### The gnomAD browser

*Support for multiple reference genomes*

Alongside the release of gnomAD v3, we wanted to retain information from previous releases (gnomAD v2) in the browser for reproducibility. To do so, we added support for multiple reference genomes to the browser (Supplementary Figure 6).

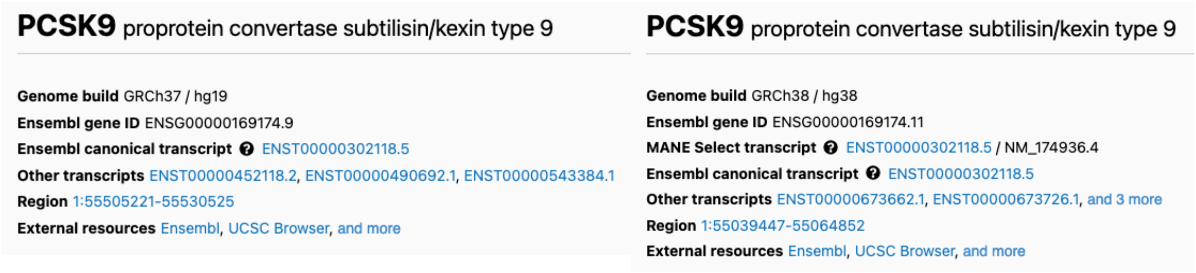


**Supplementary Figure 10 | The same gene viewed in gnomAD v2 and v3.**

Additionally, to make it easier to transition between reference builds and gnomAD versions, we added a liftover function for all variants in gnomAD v2 (Supplementary Figure 7).

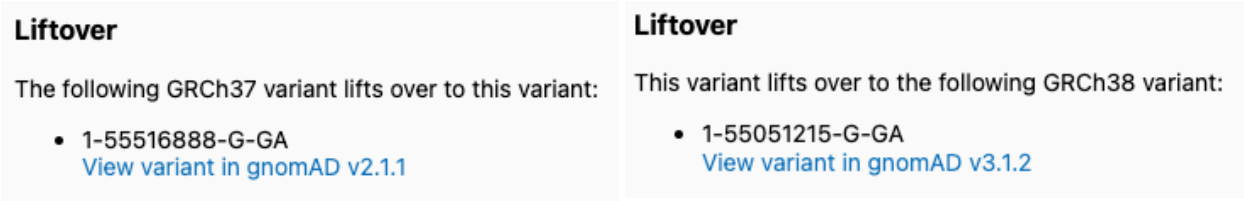


**Supplementary Figure 11 | Liftover section on a gnomAD v2 and v3 variant page.**

*HGDP and 1000 Genomes population frequencies*

For variants found in the HGDP / 1000 Genomes subset, the browser now includes population frequencies based on known populations from the HGDP / 1000 Genomes sample metadata (Supplementary Figure 8).

**Population Frequencies**

gnomAD | HGDP | 1KG

| Population | | Allele Count | Allele Number | Number of Homozygotes | Allele Frequency |
|---|---|---|---|---|---|
| European | | 21 | 1034 | 0 | 0.02031 |
| Admixed American | Overall | 11 | 686 | 0 | 0.01603 |
| | Puerto Ricans from Puerto Rico | 6 | 198 | 0 | 0.03030 |
| | Mexican Ancestry from Los Angeles, USA | 2 | 126 | 0 | 0.01587 |
| | Peruvians from Lima, Peru | 2 | 172 | 0 | 0.01163 |
| | Colombians from Medellin, Colombia | 1 | 190 | 0 | 0.005263 |
| | XX | 8 | 346 | 0 | 0.02312 |
| | XY | 3 | 340 | 0 | 0.008824 |
| African | | 0 | 1264 | 0 | 0.000 |
| East Asian | | 0 | 1002 | 0 | 0.000 |
| South Asian | | 0 | 1014 | 0 | 0.000 |
| XX | | 21 | 2506 | 0 | 0.008380 |
| XY | | 11 | 2494 | 0 | 0.004411 |
| **Total** | | **32** | **5000** | **0** | **0.006400** |

**Supplementary Figure 12 | Detailed population frequencies. Here, we show the frequency table for the 1000 Genomes project.**

*Read data in non-coding regions*

In previous releases, we provide short read data for exonic variants at the bottom of the variant page to enable detailed quality assessment. In this release, we provide short read data for all variants, including those in non-coding regions.

#### Supplementary Datasets

**Supplementary Dataset 1 | Variant counts in gnomAD genomes and mutation rates.** The number of possible and observed rare (MAF≤1%) SNVs in the 76,156 gnomAD genomes, along with the estimated mutation rate ('fitted_proportion_observed') for each trinucleotide context, reference, and alternate allele, stratified by methylation levels for CpG transitions.

**Supplementary Dataset 2 | Genome-wide constraint Z scores at 1kb scale.** A .bed file containing constraint Z scores for 1,984,900 1kb autosomal windows (passing all quality controls). Coordinates are on GRCh38.

**Supplementary Dataset 3 | Genome-wide constraint Z scores at 1kb sliding by 100bp scale.** A .bed file containing constraint Z scores for 19,834,726 1kb sliding windows (passing all quality controls). Coordinates are on GRCh38.

**Supplementary Dataset 4 | Functionally informed fine-mapping results using constraint Z as a prior.** The increase in posterior inclusion probability (PIP) when incorporating constraint Z score as a functional prior into previous fine-mapping results (that used a uniform prior; denoted as $PIP_Z$ and $PIP_{unif}$, respectively) are listed for 13,069 variant-trait pairs.

**Supplementary Dataset 5 | Constraint Z scores of enhancers linked to specific genes.** Enhancer-gene links were obtained from the Roadmap Epigenomics Enhancer-Gene Linking database. For each gene, the enhancer with the highest Z score was selected for analysis, and the membership of each gene in gene lists analyzed in Fig. 5b is annotated.

Diaz-Papkovich, Alex, Luke Anderson-Trocme, and Simon Gravel. 2018. “Revealing Multi-Scale Population Structure in Large Cohorts.” *bioRxiv*, September, 423632.

Hail Team. Hail 0.2.62-84fa81b9ea3d. https://github.com/hail-is/hail/commit/84fa81b9ea3d.

Karczewski, Konrad J., Laurent C. Francioli, Grace Tiao, Beryl B. Cummings, Jessica Alföldi, Qingbo Wang, Ryan L. Collins, et al. 2020. “The Mutational Constraint Spectrum Quantified from Variation in 141,456 Humans.” *Nature* 581 (7809): 434–43.

Karczewski, Konrad J., Matthew Solomonson, Katherine R. Chao, Julia K. Goodrich, Grace Tiao, Wenhan Lu, Bridget M. Riley-Gillis, et al. 2021. “Systematic Single-Variant and Gene-Based Association Testing of 3,700 Phenotypes in 281,850 UK Biobank Exomes.” *bioRxiv*. medRxiv. https://doi.org/10.1101/2021.06.19.21259117.

McInnes, Leland, John Healy, Nathaniel Saul, and Lukas Großberger. 2018. “UMAP: Uniform Manifold Approximation and Projection.” *Journal of Open Source Software* 3 (29): 861.

Purcell, Shaun M., Jennifer L. Moran, Menachem Fromer, Douglas Ruderfer, Nadia Solovieff, Panos Roussos, Colm O’Dushlaine, et al. 2014. “A Polygenic Burden of Rare Disruptive Mutations in Schizophrenia.” *Nature* 506 (7487): 185–90.
